# Geographic variation in the genetic basis of resistance to leaf rust between locally adapted ecotypes of the biofuel crop switchgrass (*Panicum virgatum*)

**DOI:** 10.1101/619148

**Authors:** Acer VanWallendael, Jason Bonnette, Thomas E. Juenger, Felix B. Fritschi, Philip A. Fay, Robert B. Mitchell, John Lloyd-Reilley, Francis M. Rouquette, Gary C. Bergstrom, David B. Lowry

## Abstract

- Local adaptation is an important process in plant evolution, which can be impacted by differential pathogen pressures along environmental gradients. However, the degree to which pathogen resistance loci vary in effect across space and time is incompletely described.
- To understand how the genetic architecture of resistance varies across time and geographic space, we quantified rust (*Puccinia spp.*) severity in switchgrass (*Panicum virgatum*) plantings at eight locations across the central United States for three years and conducted quantitative trait locus (QTL) mapping for rust progression.
- We mapped several variable QTLs, but two large-effect QTLs which we have named *Prr1* and *Prr2* were consistently associated with rust severity in multiple sites and years, particularly in northern sites. In contrast, there were numerous small-effect QTLs at southern sites, indicating a genotype-by-environment interaction in rust resistance loci. Interestingly, *Prr1* and *Prr2* had a strong epistatic interaction, which also varied in strength and direction of effect across space.
- Our results suggest that abiotic factors covarying with latitude interact with the genetic loci underlying plant resistance to control rust infection severity. Further, our results indicate that segregating genetic variation in epistatically interacting loci may play a key role in determining response to infection across geographic space.

## Introduction

Understanding the factors that determine how well a particular population is adapted to its environment is a major goal of evolutionary biology. Plant populations often exhibit local adaptation (Leimu & Fisher, 2008; Hereford, 2009), in which genotypes in a particular area are more successful in their local environment than foreign genotypes transplanted to that area (Kawecki & Ebert, 2004). A major component of adaptation to a local environment is adaptation to biotic factors such as competition, mutualism, herbivory, and pathogens (McKay *et al*., 2003; Fournier-Level *et al*., 2011; Price *et al*., 2018; He *et al*., 2018; Lowry *et al*., 2019). Pathogens in particular impose strong selection on plant populations and may thus influence the evolution of those populations (Giraud *et al*., 2017; Mursinoff & Tack, 2017). Disease resistance alleles vary in frequency across populations (Thrall & Burdon, 2002; Kniskern & Rausher, 2006; Chappell & Rausher, 2016) and may be differentially expressed depending on environmental conditions (Colhoun, 1973; Atkinson & Urwin, 2012; Huot *et al*., 2017). Thus, a complete understanding of local adaptation in plants requires measurement of the changes in the genetic architecture of pathogen resistance across multiple environments (Busby *et al*., 2014).

Population heterogeneity in pathogen resistance loci results in imperfect resistance for plant species. To explain maintenance of this variation, previous studies have noted the role of the abiotic environment in influencing the plant-pathogen relationship (Atkinson & Urwin 2012; Huot *et al*., 2017) as well as coevolutionary dynamics between plants and their pathogens (Thrall *et al*., 2012; Bever *et al*., 2015; Penczykowski *et al*., 2016; Kniskern & Rausher 2016), including fitness costs of both virulence and resistance (Bergstrom *et al*., 2000; Tian *et al*., 2003). The abiotic environment can influence the relationship by restricting the range of pathogens that are, for instance, humidity or temperature-sensitive (Shaw & Osbourne 2011). For example, the oomycete pathogen of dry beans *Pythium spp.* is highly effective in high-moisture conditions (Soltani *et al*., 2018). Therefore, wet-adapted bean varieties must exhibit higher *Pythium* resistance (Soltani *et al*., 2018). Similarly, abiotic conditions can directly impact plant molecular immune responses, such as in the case of temperature-dependent immunity in *Arabidopsis* (Huot *et al*., 2017). In addition, coevolution between pathogens and host metapopulations can dramatically shape plant evolution, largely independently of the abiotic environment. As plants evolve immune responses, pathogens evolve ways to evade immunity, resulting in population-specific adaptations to variable pathogen regimes (Thrall *et al*., 2012; Bever *et al*., 2015; Penczykowski *et al*., 2016; Kniskern & Rausher 2016). Determining the extent to which plant-pathogen coevolution is shaped by abiotic conditions and “arms race” dynamics requires a pathosystem studied over several years and a wide range of environmental conditions.

Switchgrass (*Panicum virgatum* L.) and its obligate fungal pathogens, especially switchgrass leaf rusts (*Puccinia spp.*), are an ideal system to study long-term changes in the molecular basis of resistance. Switchgrass is a long-lived, polyploid, C4, perennial grass native to North America east of the Rocky Mountains from northern Mexico to southern Canada (Cronquist, 1963). It is a common prairie and pasture grass grown as both a forage crop and as a bioenergy feedstock (Casler, 2012; Parrish *et al*., 2012). Switchgrass has also become an important study system for understanding the causes of ecological specialization (Casler, 2012; Lowry *et al*., 2014). Switchgrass is split into two locally adapted ecotypes, upland and lowland (Morris *et al*., 2011; Lowry *et al*., 2014; Milano *et al*., 2016). The upland ecotype is more common in northern North America, has a small stature (up to 190 cm), and has limited resistance to multiple pathogens, including rust and other fungal pathogens as well as viruses (Casler, 2012; Uppalapati *et al*., 2013; Milano *et al*., 2016; Lovell *et al*., 2016; Alexander *et al*., 2017). In contrast, the southern lowland ecotype is large (up to 285 cm) and is more resistant to fungal pathogens (Casler, 2012; Uppalapati *et al*., 2013; Milano *et al*., 2016; Lovell *et al*., 2016). While the lowland ecotype produces more biomass, it also has lower freezing tolerance (Lee *et al*., 2014; Peixoto & Sage, 2016), possibly explaining the rarity of lowland ecotypes in more northern climates. Temperature, moisture regimes, and other aspects of climate often covary with latitude in the central US, so over the switchgrass range plant traits respond to north-south clines (Lowry *et al*., 2019).

Since switchgrass ecotypes differ in their susceptibility to rust infection, this host-pathogen system is useful for testing the role of local variation in rust species presence in the evolution of resistance. Switchgrass is infected with at least five species of rust (*Puccinia spp.;* Demers *et al*., 2017). Rusts are basidiomycete fungi that infect only living leaf tissue (biotrophs). As such, these fungi are thought to be extirpated from northern switchgrass populations every winter when switchgrass senesces. Reinfection may happen from the alternate host, thought to be *Euphorbia corollata* (Demers *et al*., 2017), or through wind-borne spores from already-infected plants. Rust infection is common in switchgrass stands in North America by late summer, though damage is less severe in varieties from the lowland ecotype.

While there is still little known about the natural history of switchgrass rusts, closely-related wheat rusts such as *Puccinia striiformis* have been extensively studied, and thus provide a model for thinking about switchgrass rusts. In addition to restrictions imposed by obligate biotrophy, winter minimum temperatures below −10°C may kill fungal spores, and high nighttime temperatures limit disease development (Chen, 2005). Further, the spores require high humidity to germinate, and may be spread between plants by rainfall (Chen, 2005). The geographic range of switchgrass encompasses many temperature and precipitation regimes, so direct abiotic selection likely plays an important role in rust biology. Further, the abiotic environment directly impacts plant resistance to rust. In *P. striiformis*, resistance genes such as *Yr36* are effective at high, but not low temperatures (Fu *et al*., 2009). Thus, over the geographic range of the switchgrass-rust interactions, variation in the abiotic environment may play an important role in influencing both pathogen virulence and plant defenses.

The distribution of related pathogen species can have an important role in the evolution of resistance. Resistance mechanisms can be specific to particular rust species or to specific strains of a species. Wheat varieties bred for stem rust (*Puccinia graminis* f. sp. *tritici*) resistance were overcome in recent years by a novel rust lineage known as Ug99 (Singh *et al*., 2015). Ug99 originated from the dikaryotic parasexual combination of two asexual rust lineages (Li *et al*., 2019). If rust species are geographically restricted, switchgrass may evolve local resistance mechanisms to respond to specific rusts. In contrast, if only one species is dominant, the genetic basis of resistance may be identical over a wide geographic range, or population-specific resistance will occur at the level of pathogen strain, rather than pathogen species. There is conflicting evidence about the ranges of the five switchgrass rusts. Demers *et al*., (2017) found predominantly *P. pammelii* and *P. graminicola* in the midwestern United States, and *P. novopanici* predominantly in eastern states. In contrast, Kenaley *et al*., (2018) found that most midwestern switchgrass plantings are infected with *P. novopanici*, with *P. pammelii* and *P. graminicola* showing up only rarely.

The primary goal of our study was to understand the nature of genotype-by-environment interactions underlying variation of switchgrass rust resistance across the central United States. To overcome past limitations due to environmental and temporal variation in host-pathogen relationships, we measured fungal resistance over three years at a large geographic scale. We studied quantitative trait locus (QTL) mapping populations replicated at eight sites across more than 1500 km of latitude to map QTLs for rust resistance. These populations have been previously used to study QTL x environment interactions for several morphological and phenological traits (Lowry *et al*., 2019), but biotic interactions have not been assessed prior to this study. The geographic patterns of QTL presence and strength allowed us to examine the spatial distribution of pathogen resistance expression. If QTL presence and strength was correlated with latitude of the field sites over the eight sites, we could infer that the abiotic environment played an essential role in governing the interaction. However, if resistance was site-specific, that would be evidence of rust population dynamics as being a stronger influence on plant resistance. Further, the four mapping cross genotypes allowed a chance to test the host-population-specificity of resistance. We predicted that any resistance alleles that we mapped would originate from a lowland genotype, since lowland plants are more resistant to fungal pathogens. Further, we expected that resistance alleles would explain substantial variation in other morphological traits that distinguish ecotypes, indicating that resistance genes are associated with overall local adaptation in switchgrass.

## Methods

### Development of mapping populations

To identify loci controlling variation in rust progression, we used a previously developed four-way phase-known (pseudo-testcross) mapping population derived from both upland and lowland genotypes (conceptual map in Figure 1). For full details of the development of the mapping population see (Milano *et al*., 2016). We clonally divided the outbred populations by manually splitting rhizomes at the Brackenridge Field Laboratory in Austin, TX. In May-July of 2015, the F_0_, F_1_, and F_2_ clones were potted, moved by truck, and transplanted into the field at ten sites throughout the United States (Lowry *et al*., 2019). Thus, at each site, we planted >5 clones of each of the four F_0_ genotypes, 15 individuals of each of the two F_1_ populations, and 431 individuals from the F_2_ generation. We were able to monitor pathogen changes throughout the course of this study at eight of these sites: Kingsville, TX; Austin, TX; Temple, TX; Overton, TX; Columbia, MO; Manhattan, KS; Mead, NE; and Hickory Corners, MI (Figure 1). We assigned plants randomly to a honeycomb design, with 1.56 m between each plant. To reduce edge effects, we planted a border of lowland plants around the plot that were not measured experimentally. We watered plants by hand in 2015, when necessary to facilitate establishment. Weed cloth was installed to cover the ground between plants and reduce weed pressure. After 2015, we removed weeds using pre-emergent herbicides and physical pulling but did not otherwise manage plots. To develop a linkage map for QTL mapping we genotyped 431 second-generation genotypes by whole genome resequencing (for full sequencing details, see Lowry *et al*., 2019).

**Figure 1:**
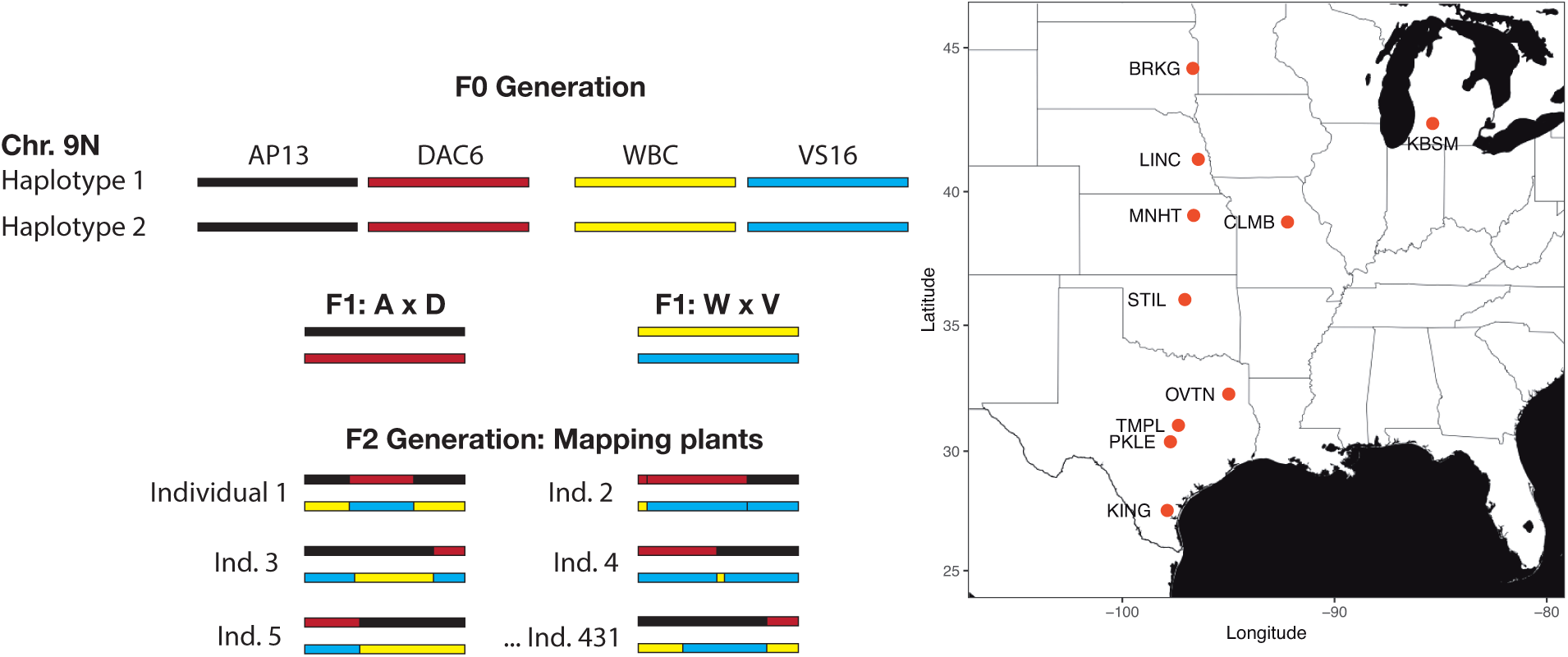
Experimental design. **a**. Conceptual map of cross design. Horizontal bars represent copies of chromosome 9N, for example. In the F0 generation (grandparents), each genotype has two mostly homozygous haplotypes, shown by the color of the bars. In the F1 cross generation, there are two combinations of haplotypes: A x D for AP13 x DAC6, and W x V for WBC3 x VS16. The F2 generation combines the F1 haplotypes, so each of the 431 offspring individuals have some combination of the four grandparental alleles at any one locus. B. Locations of experimental sites in central North America. KING: Kingsville, TX; PKLE: Austin, TX; TMPL: Temple, TX; OVTN: Overton, TX; CLMB: Columbia, MO; MNHT: Manhattan, KS; LINC: Mead, NE; KBSM: Hickory Corners, MI. We additionally collected rust samples from sites in Stillwater, OK (STIL), and Brookings, SD (BRKG), but were not able to assess rust infection scores at these sites.

### Phenotyping

At each site we scored the presence of leaf rust in 2016, 2017, and 2018. We used a method developed for rust on wheat (McNeal *et al*., 1971; Roelfs *et al*., 1992), which translates well to switchgrass and has been used in previous studies (Uppalapati *et al*., 2013). At each site, we scored rust on a 0 - 10 scale based on the total proportion of the canopy covered in rust pustules, a score which we have defined as ‘rust severity’ for this study (Figure 2). The 0 – 10 scale can be thought of as 1/10 of the percent of leaf area covered in rust pustules (a score of 1 corresponds to 10% leaf cover, 3 to 30% cover, and so on). We assessed the whole canopy visually. This subjective rating is imperfect since it relies on field ratings by technicians rather than a quantitative measure. However, alternative methods such as leaf collection and scanning are much more labor intensive and introduce similar biases when choosing leaves. Other fungal pathogens such as anthracnose and *Bipolaris* were present in plots but were much less common than rust and were not reflected in our ratings.

**Figure 2:**
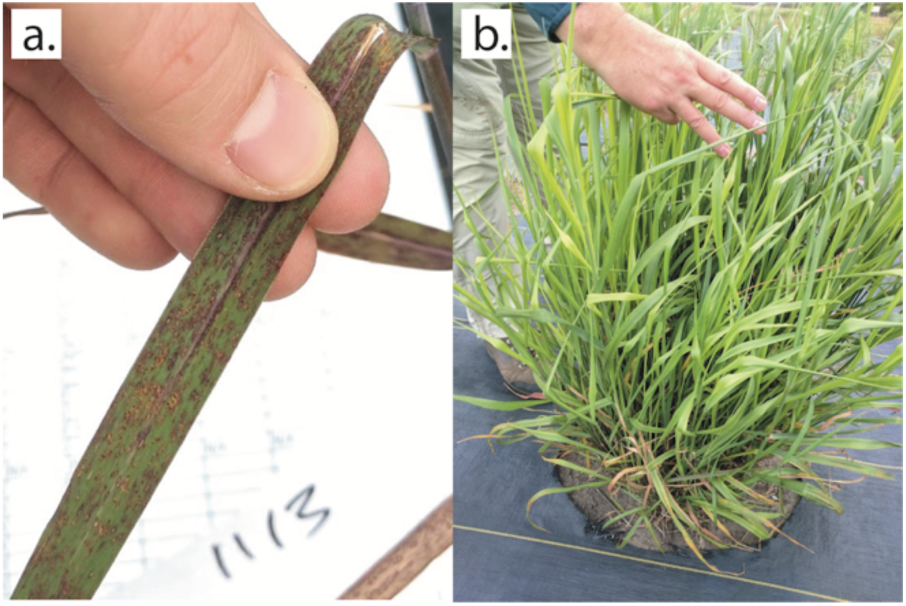
**a**. Heavily infected single leaf. **b**. Lightly infected small plant. This individual would be rated a 3, since approximately 30% of the leaf surface is covered with rust pustules.

The effort extended to phenotyping rust severity varied among sites and years due to logistical challenges. However, for the most part, sampling began three weeks after green-up (the point at which ∼50% of plants tiller crop had emerged from the soil) and continued weekly until severity stopped increasing (Figure 3). Over three years, this resulted in more than 149,000 rust ratings, which we used for the QTL analyses. In addition, we measured other morphological and physiological traits at all sites, including the number of tillers, plant height at end of season, date of first flowering, and end-of-season aboveground biomass (see Milano *et al*., 2016 and Lowry *et al*., 2019 for details of this phenotyping effort).

**Figure 3:**
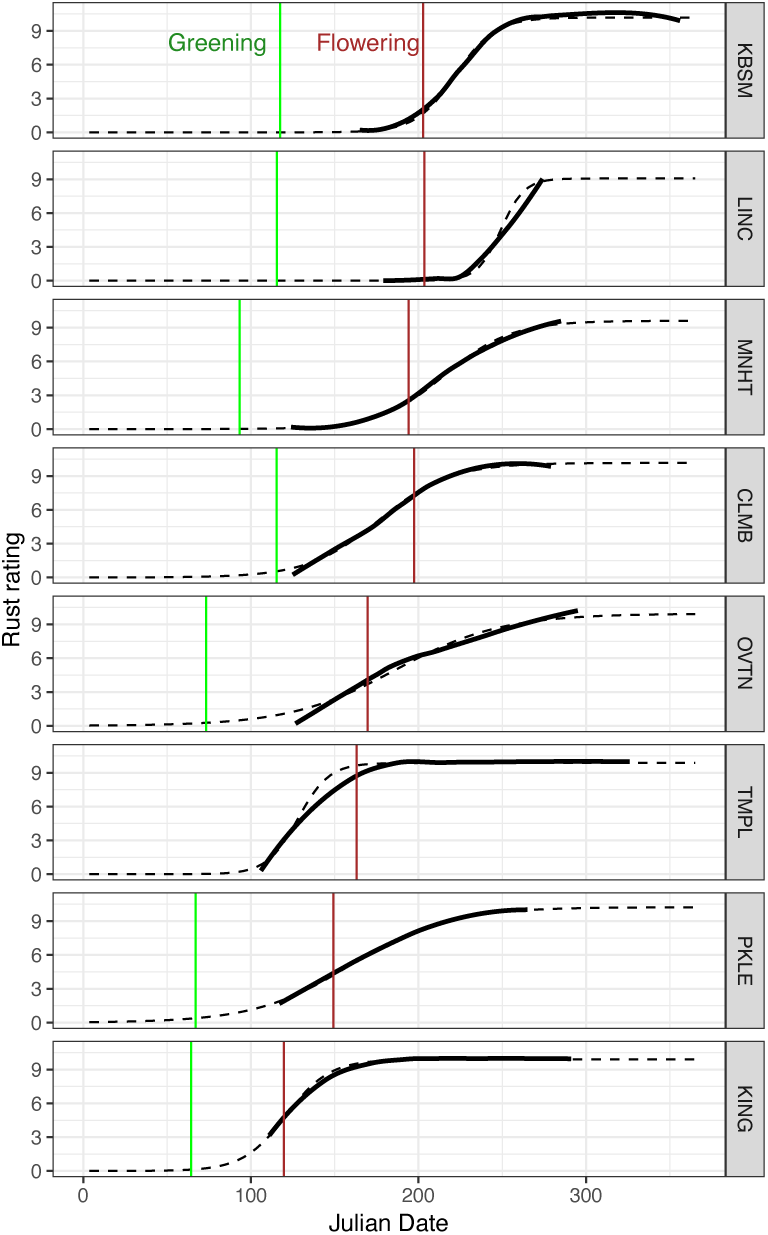
Rust progression curves in 2016. Black lines show smoothed mean rust values for sampled dates, black dotted lines show fitted logistic curves to sampled data. Green and brown vertical lines show green-up date and date of first flowering, respectively. Green-up was not quantified at TMPL in 2016 (Temple, TX; see 2017 & 2018 in Figure S4)

### Rust abundance and species composition

Visual pathogen scoring regimes are well known to be subject to statistical artifacts (Lesaffre *et al*., 2012) and our data were no exception. Pathogen scores followed a tail-inflated (“U”-shaped) distribution. Therefore, we used nonparametric Wilcoxon signed-rank and Kruskal-Wallis tests through the functions *wilcox.test* and *kruskal.test* in the *stats* package of R to test the differences in severity between lineages, years, and sites (R core team, 2018). To test for cytoplasmic effects on rust severity, we compared rust scores between F_2_ individuals with maternal cytoplasm from upland and lowland grandparents, also with the aforementioned nonparametric tests.

We assayed the prevalence of various species of rust at study sites in 2018 by sequencing a portion of the ITS (Internal Transcribed Spacer) region for subsampled isolates from each site. At each location, we haphazardly collected leaves from ∼20 individuals, and isolated single pustules from each. By sampling 20 individuals, we ensured that we would have the power to detect species present in a substantial portion of the field (>30%) at a 99.9% confidence level. We followed the same method as Kenaley *et al*., (2018) for DNA extraction, PCR amplification, purification, and sequencing. Briefly, we extracted genomic DNA from leaf segments cut from areas with one pustule or very few pustules using the DNeasy Plant Mini Kit (Qiagen, Valencia, CA). We then amplified the ITS2 region using the primer pair RUST2inv (5′-GATGAAGAACACAGTGAAA; Aime 2006) and RUST1 (5′-AGTGCACTTTATTGTGGCTCGA; Kropp *et al*., 1995), producing a 510 bp fragment. We purified the PCR product using Qiagen QIAquick PCR Purification Kit (Qiagen, Germantown, MD) and then sequenced unidirectionally with 180 ng of template, 25 picomoles of primer RUST1, and using a BigDye terminator sequencing kit (Applied Biosystems, Foster City, CA) on an ABI 3730 DNA Sequencer (Applied Biosystems).

### Genomic architecture

Initially, we mapped QTLs based on pathogen ratings for each individual time point at which rust prevalence was quantified. Pathogen ratings were processed in R (v3.4; R core team, 2018) using both packages *qtl* and *funqtl* (Broman *et al*., 2003; Kwak *et al*., 2016). To examine QTL effects over time, we scanned for QTLs using Haley-Knott regression for each site by year combination, with each time point as a separate trait using the functions *scanone* in *qtl* and *geteffects* in *funqtl* (Broman *et al*., 2003; Kwak *et al*., 2016).

We additionally examined QTLs controlling the overall progression of rust by modeling severity as a function-valued trait (Kwak *et al*., 2014). For function-valued traits such as time-series, QTLs can be summarized across time using the R package *funqtl* (Kwak *et al*., 2014; 2016). This method has the advantage of decreasing bias introduced by differences among raters at different sites as well as phenology differences by summarizing multiple QTLs using a penalized likelihood approach (Kwak et al. 2014). Previous studies have used area under disease progress curve (AUDPC) measurements to quantify resistance (Jeger & Viljanen-Rollinson, 2001). AUDPC is robust and useful but may show bias when infection timing differs among sites (Jeger & Viljanen-Rollinson, 2001). The *funqtl* method yields two scores for each phenotype, the mean LOD (SLOD), and the maximum LOD (MLOD). The first is useful for identifying loci that show large effects over time, while the second identifies loci that have a large effect for a single time point (Kwak et al. 2014). Since we were most interested in QTLs with effects across the season, we focused on SLOD scores. We conducted 1000 permutations to calculate a penalty for the SLOD score that reduces the rate of inclusion of extra loci to 5% (Broman & Sen, 2009). To estimate the percentage explained variation (PEV) for each significant QTL, we fit single-QTL models for each time point with the *fitqtl* command in the *qtl* package, then used the maximum PEV value (Broman *et al*., 2013). We examined geographic and temporal variation by mapping QTLs separately for each site and year, but we also generated a combined test that summarizes variation in this experiment. For this test, we summed SLOD scores across multiple sites and years, and concatenated permutations to generate a critical SLOD cutoff. We produced all plots using *funqtl* and *ggplot2* (Kwak *et al*., 2016; Wickham, 2016).

Finally, we estimated the allele-specific effects of the significant QTLs we discovered. In the four-way cross design, second-generation offspring will have one of four possible genotypes at each locus: AP13 & WBC3, AP13 & VS16, WBC3 & DAC6, or DAC6 & VS16. These genotypes represent the combination of alleles from each of the four grandparents of the cross (lowland AP13 and WBC; upland DAC and VS16; Figure 1). For each locus, we compared the pathogen scores for the individuals with the “resistant” QTL alleles to those with the “susceptible” alleles. We additionally made this comparison for morphological traits, including biomass, flowering time, tiller count, and green-up date. We tested for difference in means using nonparametric Wilcoxon signed-rank tests for pathogen ratings through function *wilcox.test* in the *stats* package, and a two-sample *t*-test for all morphological traits through function *t.test* in the *stats* package (R core team, 2019). R code for all analyses is available for download on the author’s Github page: https://github.com/avanwallendael/switchgrass_rust_qtl.

## Results

### Rust abundance and composition

Though infection timing varied, rust was present at all sites throughout the study period (Figure 3). The largest divergence in infection was between upland and lowland F_0_ (grandparental) plants, with upland plants experiencing 39.15% more rust than lowland plants (*W* = 1.15e7, *P <* 0.0001). The divergence between lowland and upland plants in rust severity was comparable in effect size at all sites, and never differed in direction. Rust severity also differed between generations in the cross (*χ^2^* = 656.98, *P <* 0.0001), with F_0_ plants showing the least amount of rust and F_1_ plants the greatest (Figure S1). Rust severity was negatively correlated with green-up date, biomass, flowering time, height, and tiller count (Figure S2). Field sites differed substantially in mean rust severity across years (*χ ^2^* = 1.99 x 10^-5^, *P <* 0.0001; Figure S3), though this was confounded by different sampling periods across sites. Mean rust severity declined in almost every site each year (Figure S3), decreasing by 19.75% in 2017 and an additional 30.74% in 2018. This change correlates with biomass increases of 85.64% in 2017 and an additional 46.89% in 2018. Our cross design allowed us to test the phenotypic effect of maternal cytoplasm, the difference between second-generation plants with an upland seed parent and those with a lowland seed parent. We compared rust scores between second-generation individuals with maternal cytoplasm from upland and lowland F_1_s. There was a significant cytoplasmic effect (*W* = 1.72e9, *P =* 0.0059), but rust scores were only 1.33% higher in plants with lowland (WBC) cytoplasm than plants with the upland (DAC) cytoplasm (Figure S1).

We assayed the prevalence of various species of rust at the study sites in 2018 by sequencing ITS2 (Internal Transcribed Spacer 2) for subsampled isolates from each site. While we found evidence of three rust species infecting switchgrass, *Puccinia novopanici* was the predominant rust species at all field sites that we examined (Table 1). *P. graminicola* was present only in Brookings, SD (BRKG) and Mead, NE (LINC), while *P. cumminsii* was present in only one sample from Austin, TX (PKLE). Given the rarity of other *Puccinia spp.*, we did not detect statistical differences between genotypes or sites in pathogen species composition, with the exception of Mead, NE (LINC), which was 35% *P. graminicola*.

**Table 1:**
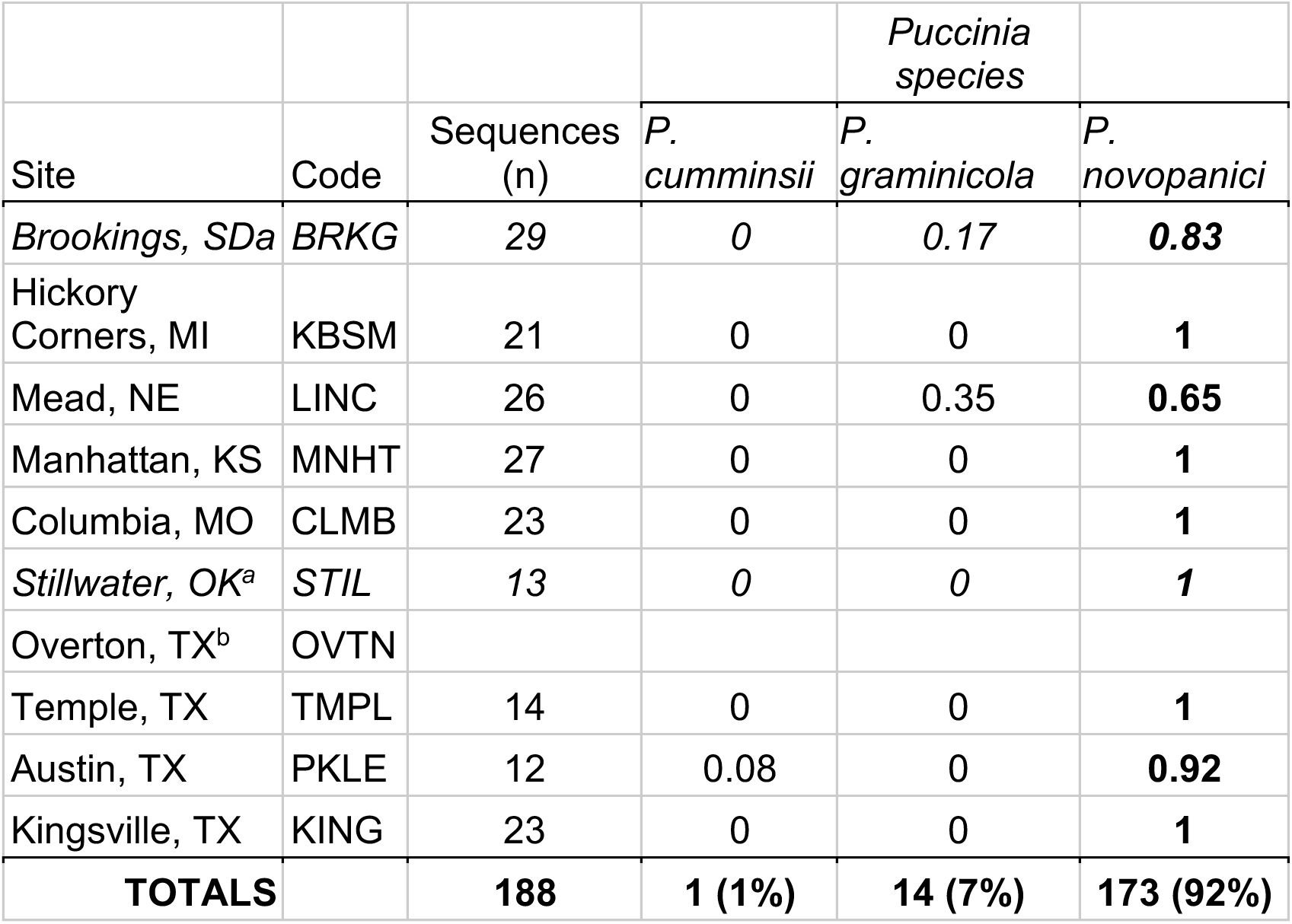
Species of rust detected by field sites, with values shown as the proportion of viable ITS sequences that matched each species. ^a^STIL & BRKG were not used in QTL mapping due to missing rust severity measurements. ^b^We were not able to obtain samples from OVTN for <u>species identification.</u>

### Genomic architecture

Since we collected data on pathogens over several weeks at all sites, we were able to examine how the genomic architecture of resistance changed over a single season at each site by treating each time point as a distinct phenotype for QTL mapping. For simplicity throughout this paper, we refer to rust resistance as: Resistance = (1 – Severity) (Simms & Triplett, 1994). We found extensive variation between sites, but some patterns were shared across several sites. We identified QTLs for resistance on chromosomes 3N and 9N that were consistently associated with rust resistance, which we have named loci *Prr1* (*PUCCINIA* RUST RESISTANT 1) and *Prr2* (*PUCCINIA* RUST RESISTANT 2), respectively. The lowland allele (AP13) only conferred greater resistance for one of these loci (*Prr2*), relative to the upland allele (DAC). In contrast, the lowland allele (AP13) increased rust severity relative to the upland allele (DAC) on chromosome 3N (*Prr1*). Over the course of the field season, these QTLs showed effects for ∼40-50 days when mapped for single time point measurements, with *Prr1* becoming detectable about one week after *Prr2* (Figure 4 and Supp.). Overall patterns were similar across years, although the collection of fewer time points in 2017 resulted in lower-resolution data.

**Figure 4:**
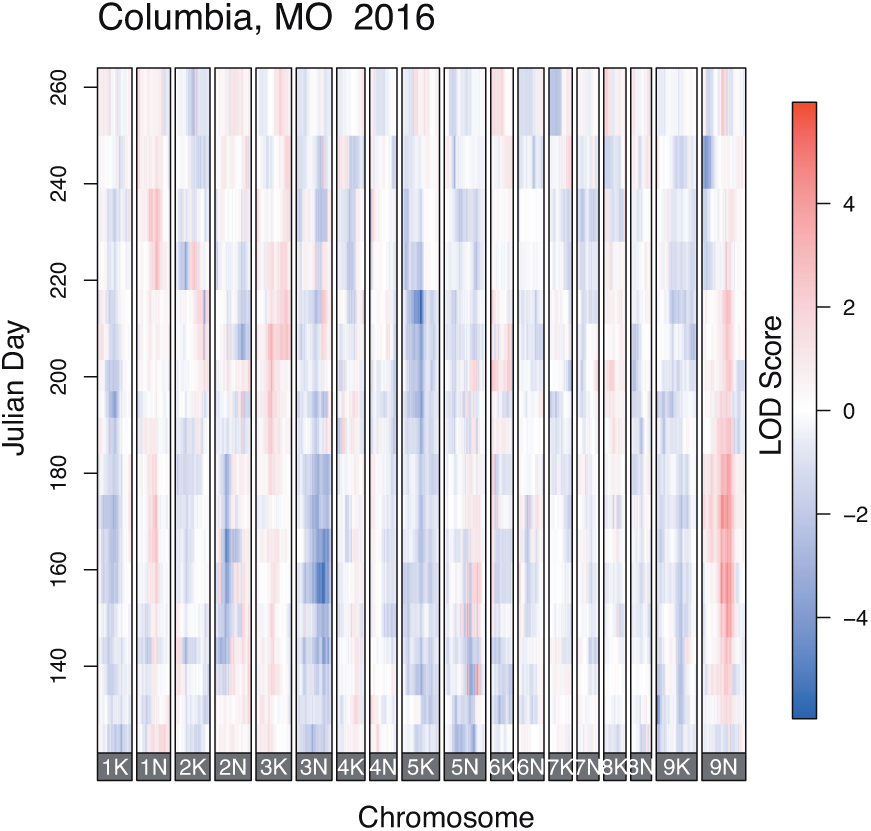
Time-series QTL effects for Columbia, MO in 2016 (remaining sites & years in Figure S5). Blue indicates that the lowland allele decreases rust, red indicates that the upland allele decreases rust. Color intensity is proportional to LOD score, or the strength of the QTL.

To evaluate the genetic architecture of resistance over the entire field seasons, we also mapped QTLs to rust severity as a function-valued trait. Summing across all sites and years, we found 51 total significant QTLs (Figure 5a; Table S1). These QTLs varied in percent of explained variation (PEV) throughout the infection season, with a single QTL model explaining up to 24.2% of the variation in rust severity (Table S1). Many of these QTLs were at overlapping genomic positions, so overall, we found 18 locations throughout the switchgrass genome that were associated with rust in at least one site. Just 4 of the 18 switchgrass chromosomes exhibited no QTLs at any position. Overall, we found the highest number of QTLs at the most northern site (KBSM), though there was not a clear geographic pattern in QTL number. Additionally, there was variation between years, with the greatest number of QTLs in 2016 (24 QTLs), and fewer in 2017 and 2018 (15 and 12, respectively). When we combined QTLs across sites and years (Figure 5C), *Prr1* and *Prr2* had the highest LOD scores in the north.

**Figure 5:**
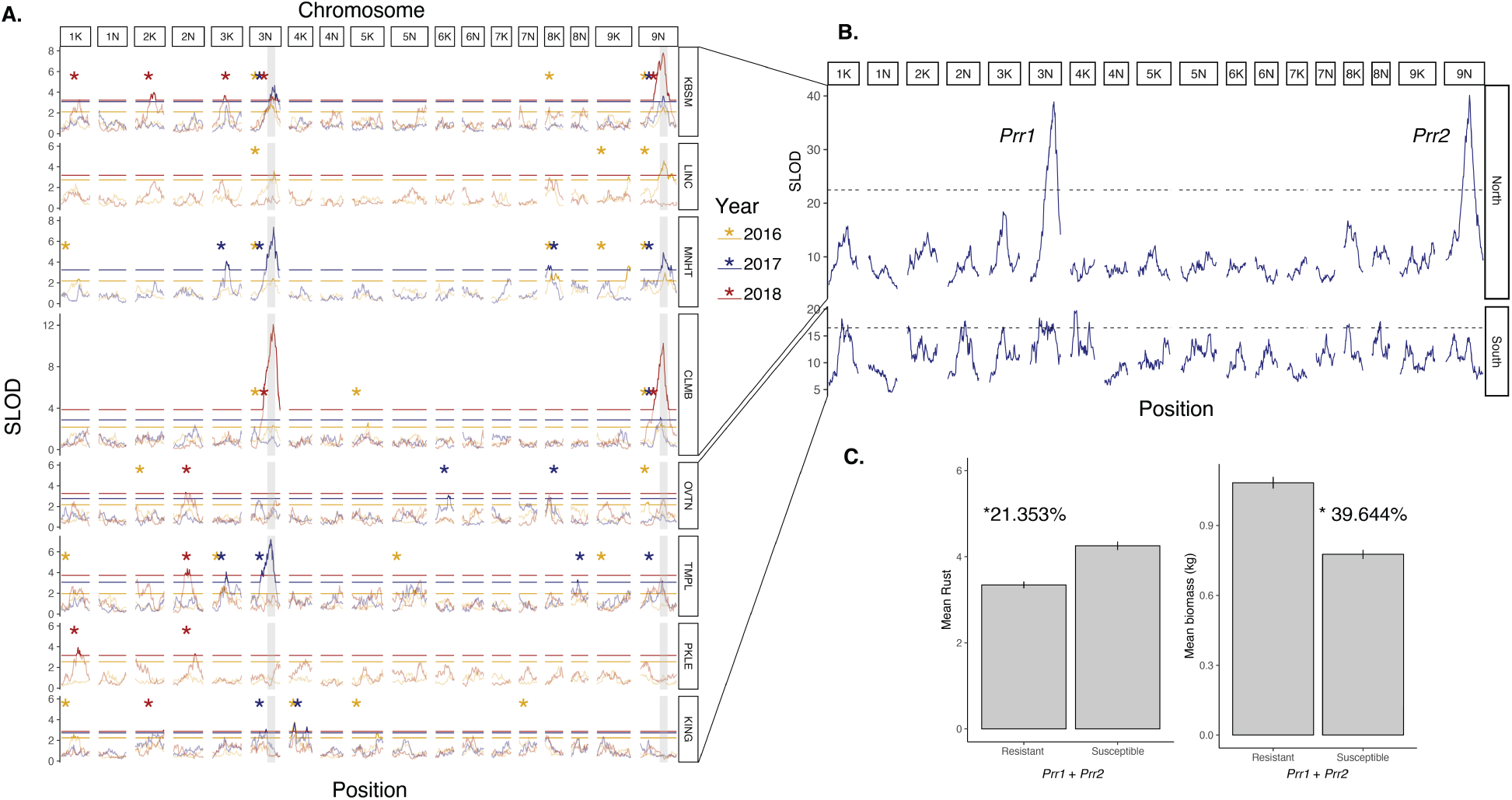
**a**: QTLs for all sites and years. SLOD shows the mean log-odds score for function-valued traits, or the strength of the QTL. Horizontal lines show significance thresholds for each test, peaks above the threshold are shown as solid lines, and peaks below as faded lines. Vertical gray bars highlight large-effect QTLs *Prr1* & *Prr2*. Asterisks show chromosomes that had at least one significant QTL and are colored by year. **b**: Combined SLOD plot showing SLOD scores summed over Northern and Southern sites. **c**: The phenotypic effects of *Prr1* & *Prr2* on rust infection in the early summer season and total yearly biomass.

Given their strength and consistency across sites and years, we considered *Prr2* and *Prr1* as the most important QTLs. These large-effect QTLs differed greatly in their frequency of significant effect between northern sites and southern sites, showing significant effects in northern four sites 77.2% (17/22) of the time over the three years, but only 16.6% (4/24) of the time in southern four sites. When the southern sites were pooled, we detected nine significant QTLs across the southern sites, but these were much weaker in effect (Figure 5b).

### Allelic effects of Prr1 and Prr2

We calculated the combined effects of *Prr1* and *Prr2* by examining only individuals containing either resistant or susceptible alleles at each locus. For instance, the AP13 allele increased resistance at *Prr1*, but the DAC6 allele increased resistance at *Prr2*, so the individuals with both of these alleles were designated as having the “resistant” combination. We compared these resistant plants to those that had susceptible alleles at both *Prr1* and *Prr2* to assess the combined effects of the QTLs.

The effects of *Prr1* and *Prr2* were clear in their impacts on rust severity. The combination of “resistant” alleles across loci resulted in 21.35% lower rust scores across the season (*W* = 31808000, *P <* 0.0001; Figure 5B, Table 2). In Southern sites, there was no clear difference between “resistant” and “susceptible” alleles’ impact on rust (*W* = 734, *P* = 0.34). To establish whether *Prr2* and *Prr1* also impacted other important phenotypes, we conducted additional contrasts between the resistant and susceptible alleles. The resistance alleles were associated with 39.6% higher biomass (*t* = 12.34, *P <* 0.0001, Figure 5), 17.36% greater tiller count (*t* = 6.71, *P <* 0.0001), and 5.99% greater height (*t* = 7.49, *P <* 0.0001), but had little effect on flowering time (*t* = 1.94, *P =* 0.052; Table 2).

**Table 2:**
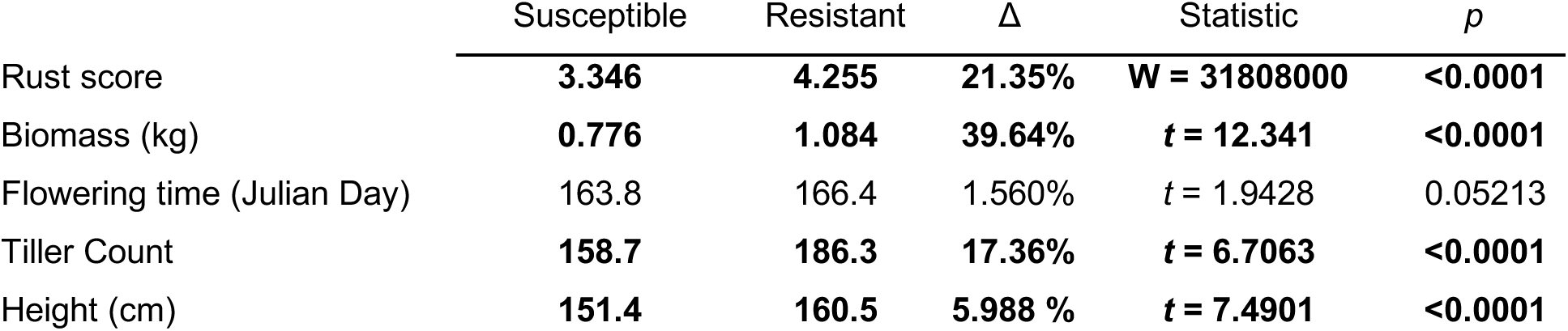
Genotype-specific effects for the combination of *Prr1* and *Prr2* for rust resistance, with bold values significant at α = 0.05.

Additionally, we performed a simple test for epistasis between large-effect loci. Since performing epistasis tests for all QTLs, phenotypes, sites, and years would be computationally prohibitive, we tested one phenotype (peak rust) per site and year for the two large-effect loci only. We fit with a linear model (ANOVA) to test for epistasis among these two loci: *R = Prr1 + Prr2 + Prr1*Prr2 + e*, where *R* is the rust severity, *Prr1* and *Prr2* are the genotypes at those loci, and *e* is the residual error term. Epistasis is represented by a significant interaction term at a Bonferroni-corrected α of 0.00263 (α = 0.05 / 19 tests). We used the functions *anova* and *lm* in the *stats* package of R and tested for allele-specific effects using orthogonal contrasts (R core team, 2019).

We found strong evidence for an epistatic interaction between *Prr1* and *Prr2* (Figure 6) that impacted both rust severity and overall biomass. For each site and year that both QTLs were present, the interaction was significant (Figure 6; Table S2). The interaction was largest in Columbia, MO in 2018, where the effect of both QTLs was the strongest (F = 29.7, *P <* 0.0001; Figure 6C-D). This effect was driven by the combination of upland and lowland alleles inducing rust susceptibility in the mapping population. Individuals with the DAC6 allele at *Prr1* and the AP13 allele at *Prr2* were on average 10.6 times more susceptible to rust damage (t = 16.108, P < 0.0001). In addition, the allele combinations that increase rust tend to also result in lower biomass (Figure 6D). However, infrequently the direction of the rust epistatic interaction was reversed, but the biomass interaction was not (Figure 6E-F). At the KBSM site in 2018, individuals with the DAC6 allele at *Prr1* and the AP13 allele at *Prr2* were on average 2.28 times more resistant to rust damage (*t* = 9.560, *P* < 0.0001). This pattern was also observed to a lesser degree in KBSM in 2016 and TMPL in 2018.

**Figure 6:**
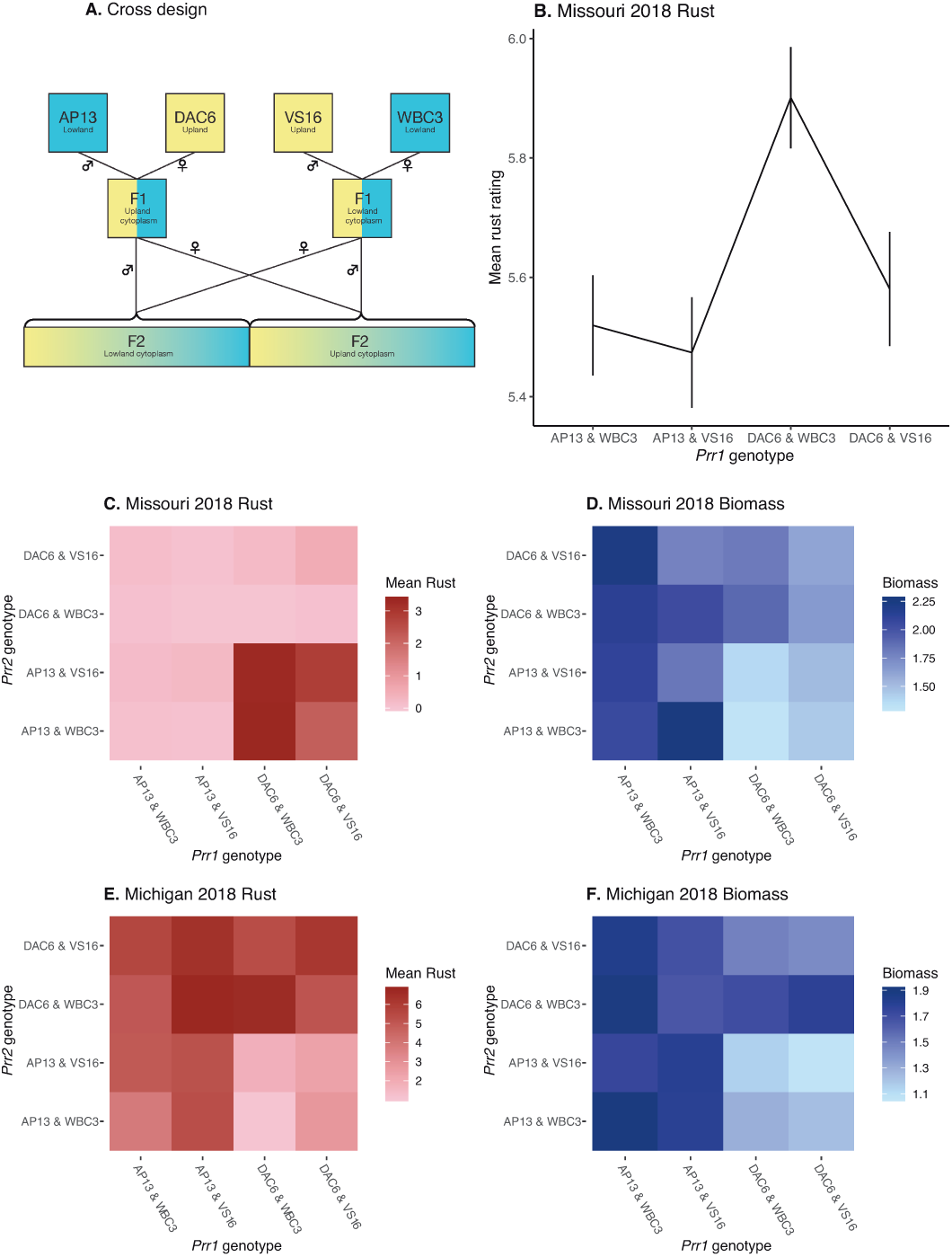
Pattern of genotype-specific effects for focal QTLs on chromosomes 3N and 9N. **A:** Four-way cross design. Two lowland genotypes, AP13 and WBC3, and two upland genotypes, DAC6 and WBC3 were crossed to create two F_1_ populations. The F_1_ populations were then crossed to create F_2_ populations. **B:** An example of phenotypic effects at a single QTL. At the 3N QTL, each individual in the F_2_ population has one of four combinations of F0 alleles, shown on the x-axis. The average effect of those genotypes on rust severity can be measured as a mean of the ratings for all F2 individuals with that genotypes at that locus. **C-F:** Epistatic interactions between specific genotypes at the 3N (*Prr1*) and 9N (*Prr2*) QTLs for both rust severity (red hues) and biomass (blue hues) at two different sites in 2018.

## Discussion

Overall, we found that there is a marked difference in the genetic architecture of rust resistance in the northern versus southern regions of the central United States. The two large-effect QTLs, *Prr1* and *Prr2*, consistently explained variation in rust severity in the north. In contrast, these large-effect QTLs rarely had significant effects in southern sites, indicating that the genetic architecture of rust resistance is highly dependent on environmental context. The effects of *Prr1* and *Prr2* were largely stable across all three years of the study, indicating that they confer consistent, but region-specific, resistance. *Prr1* and *Prr2* also co-localize with QTLs for biomass, tiller count, height, and flowering time. In contrast to our prediction that the lowland grandparental alleles would provide the most rust resistance, we found that the upland allele at *Prr1* increases resistance. An epistatic interaction between upland (DAC) and lowland alleles (AP13) at *Prr1* and *Prr2* causes greatly increased rust susceptibility in most sites and years, but this interaction was reversed for some sites across different years of the study. We discuss the implications of these results below.

### North-South difference in QTL expression

The great difference in the genetic architecture of resistance between southern and northern field sites suggests that variation in climate, correlated with latitude, may directly impact the expression of resistance to switchgrass rust. We anticipated that variation across field sites in QTL effects could be caused by differences in rust species composition at each of these sites. However, we found an overwhelming predominance of one species, *Puccinia novopanici*, at all sites other than Mead, NE (LINC). While we were not able to sample species composition for all years of this study, the dominance of *P. novopanici* in the central United States is corroborated by previous reports on this system (Kenaley *et al*., 2018). The hypothesis that large-scale climatic variation accounts for QTL variation in switchgrass was also supported in another study that used these same mapping populations (Lowry *et al*., 2019). That study showed evidence that many QTLs for morphological and phenological traits, including several that colocalize with *Prr1* and *Prr2*, exhibit GxE interactions across the north-south range of the experiment, and that overall climatic variation across the range explains the gradient (Lowry *et al*., 2019). Greater knowledge of the genes underlying these QTLs would improve our understanding of these interactions, but annotation in the switchgrass genome is incomplete as of the time of this publication. Eleven genes are in the 1.5 LOD-drop interval for *Prr1*, and forty-eight in the interval for *Prr2*, but none contain clear immune motifs such as leucine-rich repeat regions, or are known to be part of established immune defense pathways (list of genes in Table S3)

The north-south split implicates an abiotic pattern, but the means by which climate influences resistance is still undetermined. In other pathosystems, rust strain diversity and expression of resistance genes have been documented to cause variation in resistance. Pathogens often exhibit strain-specific avirulence genes that may or may not be recognized by plant R-genes (Jones & Dangl 2006). Therefore, over geographic space, the distribution of pathogen genetic diversity can determine host resistance expression (Chappell & Rausher 2016). Rust pathogens can be geographically constrained by their low freezing tolerance and the distribution of their alternate host (Kenaley *et al*., 2018). Therefore, if rust population diversity is higher in southern regions, as is true for wheat rusts in Asia (Ali *et al*., 2014), switchgrass resistance expression may involve more loci than in northern populations, resulting in a more complicated genetic architecture.

Alternatively, the mode of resistance itself may directly respond to the climate. In Arabidopsis, a change from 10-23°C to 23-32°C is sufficient to induce a change from the specific immune system (Effector-Triggered Immunity; ETI) to the generalized immune system (Pattern-Triggered Immunity; PTI; Cheng *et al*., 2013). Wheat rust defense mechanisms have been specifically documented to show temperature-dependence (Fu *et al*., 2009). The northern sites in our study were planted in either hardiness zone 5 or 6, while the southern sites were in either zone 8 or 9 (Daly *et al*., 2012), suggesting that temperature may play a role. However, we cannot be certain that resistance mechanisms are directly impacted by temperature until we can isolate the genes responsible for resistance. Further work in this system should focus on determining the genetic distributions of *P. novopanici* populations to better understand whether resistance is strain-specific, and identify potential resistance genes underlying large-effect QTLs.

### Allele-specific effects and epistatic interactions of Prr2 and Prr1

We initially expected that resistance alleles would come exclusively from lowland grandparents, since lowland genotypes are more rust-resistant in all sites we planted. However, F_2_ individuals with an upland DAC6 allele at *Prr2* were overall more resistant to rust than those with lowland AP13 allele. This indicates that upland genotypes harbor resistance alleles that diverge from lowland resistance.

Further, the presence of a negative epistatic interaction between *Prr2* and *Prr1* indicates that genes underlying these loci interact directly to impact rust resistance. Epistatic interactions are commonly found in studies of pathogen resistance in plants. For example, recent QTL studies of wheat stripe and leaf rust resistance identified multiple significant epistatic interactions (Singh *et al*., 2014; Vazquez *et al*., 2015; Zeng *et al*., 2019). Despite the commonality of epistatic interactions among pathogen resistance loci, we were surprised by the degree to which the epistatic interaction was negatively synergistic, especially at sites like CLMB in 2018. This negative epistasis between an upland allele at one locus and a lowland allele is similar to a Bateson-Dobzhansky-Muller incompatibility (BDMI), an epistatic interaction that decreases fitness when independently evolving alleles are brought together through hybridization (reviewed in Fishman & Sweigart, 2018). Pathogen resistance genes have often been implicated in BDMIs, though negative epistasis of incompatible resistance loci generally causes an autoimmune response (reviewed in Bomblies & Weigel, 2007; Traw & Bergelson, 2010; see also Atanasov *et al*., 2018). However, in switchgrass, the incompatibility usually results in lower pathogen resistance, suggesting a different mechanism.

As discussed earlier, pathogen strain diversity can play an important role in determining the spatial distribution of host resistance (Chappell & Rausher 2016). The switchgrass genotype-specificity of the epistatic interaction, as well as the reversal of the pattern in certain sites and years indicates that strain-specificity may be involved. If overall resistance between lowland and upland ecotypes were explained by generalized differences in fungal resistance genes, we would expect that resistance mechanisms would segregate by ecotype. Instead, we see that particular F_0_ alleles and allele combinations can have large impacts on rust severity. The large impact of the DAC6 – AP13 epistatic interaction may indicate that these parental populations have evolved separate and mutually exclusive resistance mechanisms.

The rare reversal of the epistatic interaction from negatively to positively synergistic also is evidence of strain-specificity. In the epistatic interaction, F_2_ individuals with the lowland AP13 allele at *Prr2* and the upland DAC6 allele at *Prr1* are typically more susceptible to the rust pathogen (Figure 6C). This susceptibility is reversed in several sites and years, however, with the lowland AP13 allele at *Prr2* and the upland DAC6 allele at *Prr1* causing resistance (Figure 6E). While it is possible that this change was driven by climate, the sites and years for which the pattern was reversed were divergent in any major component of climate. It seems more likely that a different strain of rust was more prevalent for which the *Prr1-Prr2* interaction was able to provide resistance. Greater knowledge of the genes underlying these loci will be necessary to understand the nature of this strain-specificity.

### The interaction of rust resistance interaction with other traits

Typically, resistance alleles were associated with higher biomass and other overall morphological traits. *Prr2* and *Prr1* QTLs colocalize with biomass and tiller count QTLs found in a previous study (Lowry *et al*., 2019), suggesting either close linkage or a pleiotropic effect (Figure S6). That is, these loci may contain genes for resistance that are in close genetic linkage with genes that influence biomass, or resistance may directly increase biomass and tiller count by improving the health of the plant. One piece of evidence that favors close linkage over pleiotropy is that when the epistatic pattern is reversed for rust resistance, it is not reversed for biomass (Figures 6D & 6F). This close linkage of phenotypically important traits indicates that these particular genomic regions may have been important in underlying the ecotypic divergence between lowland and upland switchgrass. Future work in this system should focus on determining the genes underlying the *Prr1* and *Prr2* QTLs to map the distribution of alternative alleles at these loci across the range of switchgrass.

We note also that rust ratings on average generally decreased over the course of this study (Figure S3), which may have been due to an increase in resistance but is more likely due to the fact that older plants grow more quickly before disease onset, and therefore have a lower proportion of canopy infection. The high correlation between switchgrass traits and strong epistasis between major resistance loci indicate that future breeding efforts will be complicated.

## Conclusion

Our results show the temporal and geographic variation in the genetic architecture of rust resistance in two locally adapted switchgrass ecotypes. Two large-effect loci explain both pathogen defense and morphological differences between ecotypes, but show a limited effectiveness in the south. We found little evidence for the possibility that this pattern is driven by pathogen species differences, since rust populations were dominated by a single species, *Puccinia novopanici*. This pattern raises important questions about the drivers of genetic architecture of pathogen resistance and underscores the importance of assaying pathogen resistance across both time and space to capture the inherent variability in the interplay of biotic and abiotic drivers of genetic change. Further, the importance of an epistatic interaction in shaping variation in resistance shows the challenges of single-gene models for pathogen resistance. Future work in this system will focus on measuring the genetic variation in the rust pathogen strains, and uncovering the genes underlying *Prr1* and *Prr2*.

## Acknowledgements

We wish to thank technicians L. Vormwald, M. Iceberg, N. Ryan, P. Duberney, L. Simon, J. Sanley, K. Barthel, and M. Carey for their tireless work collecting data. S. Kenaley, with assistance from K. Myers, J. Starr, and C. Wijewardana, was instrumental in analyzing rust species data. We thank each host university for assistance maintaining field sites. The Lowry and Juenger labs, especially A. MacQueen and R. Heckman, contributed important comments and valuable discussions. This work was supported by U.S. Department of Energy (DE-SC0014156 to TEJ and DE-SC0017883 to DBL and GCB). National Science Foundation Plant Genome Research Program Awards (IOS-0922457 and IOS-1444533) to TEJ, and supported in part by the Great Lakes Bioenergy Research Center, U.S. Department of Energy, Office of Science, Office of Biological and Environmental Research under (DE-SC0018409 and DE-FC02-07ER64494). Pathogen material was shipped under APHIS permit P526-15-03302 for GCB. USDA is an Equal Employment Opportunity provider.

## Author contributions

DBL, GCB, and TEJ designed the research. All authors contributed to performance of the research. AV, JEB, TEJ, GCB, and DBL analyzed and interpreted data. AV, JEB, and GCB collected data. AV, DBL, and TEJ wrote the manuscript.

## References

Aime MC, Matheny PB, Henk DA, Frieders EM, Nilsson RH, et al., 2006. An overview of the higher level classification of Pucciniomycotina based on combined analyses of nuclear large and small subunit rDNA sequences. Mycologia. 98:896–905.

Alexander HM, Bruns E, Schebor H, Malmstrom CM. 2017. Crop-associated virus infection in a native perennial grass: reduction in plant fitness and dynamic patterns of virus detection. J. Ecol. 105:1021–1031.

Ali S, Gladieux P, Leconte M, Gautier A, Justesen AF, et al., 2014. Origin, migration routes and worldwide population genetic structure of the wheat yellow rust pathogen *Puccinia striiformis* f. sp. *tritici*. PLoS Pathog. 10:e1003903.

Atanasov KE, Liu C, Erban A, Kopka J, Parker JE, Alcázar R. 2018. NLR mutations suppressing immune hybrid incompatibility and their effects on disease resistance. Plant Physiol. 177:1152–1169.

Atkinson NJ, Urwin PE. 2012. The interaction of plant biotic and abiotic stresses: from genes to the field. J. Exp. Bot. 63:3523–3543.

Bergstrom CT, Lipsitch M, Levin BR. 2000. Natural selection, infectious transfer and the existence conditions for bacterial plasmids. Genetics. 155:1505–1519.

Bever JD, Mangan SA, Alexander HM. 2015. Maintenance of plant species diversity by pathogens. Annu. Rev. Ecol Evol. Syst. 46:305–325.

Bomblies K, Weigel D. 2007. Hybrid necrosis: autoimmunity as a potential gene-flow barrier in plant species. Nat. Rev. Genet. 8:382–393.

Broman KW, Sen S. 2009. A Guide to QTL Mapping with R/qtl, Vol. v. Springer.

Broman KW, Wu H, Sen Ś, Churchill GA. 2003. R/qtl: QTL mapping in experimental crosses. Bioinformatics. 19:889–890.

Busby PE, Newcombe G, Dirzo R, Whitham TG. 2014. Differentiating genetic and environmental drivers of plant– pathogen community interactions. J. Ecol. 102:1300–1309.

Casler MD. 2012. Switchgrass breeding, genetics, and genomics. In Switchgrass, pp. 29–53. Springer.

Chappell TM, Rausher MD. 2016. Evolution of host range in *Coleosporium ipomoeae*, a plant pathogen with multiple hosts. Proc. Natl. Acad. Sci. 113:5346–5351.

Chen XM. 2005. Epidemiology and control of stripe rust [*Puccinia striiformis* f. sp. *tritici*] on wheat. Can. J. Plant Pathol. 27:314–337.

Cheng C, Gao X, Feng B, Sheen J, Shan L, He P. 2013. Plant immune response to pathogens differs with changing temperatures. Nat. Commun. 4:1–9.

Colhoun J. 1973. Effects of environmental factors on plant disease. Annu. Rev. Phytopathol. 11:343–364.

Daly C, Widrlechner MP, Halbleib MD, Smith JI, Gibson WP. 2012. Development of a new USDA plant hardiness zone map for the United States. J. Appl. Meteorol. Climatol. 51:242–264.

Demers JE, Liu M, Hambleton S, Castlebury LA. 2017. Rust fungi on *Panicum*. Mycologia. 109:1–17.

Fishman L, Sweigart AL. 2018. When two rights make a wrong: the evolutionary genetics of plant hybrid incompatibilities. Annu. Rev. Plant Biol. 69:707–731.

Fournier-Level A, Korte A, Cooper MD, Nordborg M, Schmitt J, Wilczek AM. 2011. A map of local adaptation in *Arabidopsis thaliana*. Science. 334:86–89.

Fu D, Uauy C, Distelfeld A, Blechl A, Epstein L, et al., 2009. A kinase-START gene confers temperature-dependent resistance to wheat stripe rust. Science. 323:1357–1360.

Giraud T, Koskella B, Laine A. 2017. Introduction: microbial local adaptation: insights from natural populations, genomics and experimental evolution. Mol. Ecol. 26:1703–1710.

He L, Wu W, Zinta G, Yang L, Wang D, et al., 2018. A naturally occurring epiallele associates with leaf senescence and local climate adaptation in *Arabidopsis* accessions. Nat. Commun. 9:1–11.

Hereford J. 2009. A quantitative survey of local adaptation and fitness trade-offs. Am. Nat. 173:579–588

Huot B, Castroverde CDM, Velásquez AC, Hubbard E, Pulman JA, et al., 2017. Dual impact of elevated temperature on plant defence and bacterial virulence in *Arabidopsis*. Nat. Commun. 8:1–12.

Jeger MJ, Viljanen-Rollinson SLH. 2001. The use of the area under the disease-progress curve (AUDPC) to assess quantitative disease resistance in crop cultivars. Theor. Appl. Genet. 102:32–40.

Jones JDG, Dangl JL. 2006. The plant immune system. Nature. 444:323–329.

Kawecki TJ, Ebert D. 2004. Conceptual issues in local adaptation. Ecol. Lett. 7:1225–1241.

Kenaley SC, Quan M, Aime MC, Bergstrom GC. 2018. New insight into the species diversity and life cycles of rust fungi (Pucciniales) affecting bioenergy switchgrass (*Panicum virgatum*) in the Eastern and Central United States. Mycol. Prog. 17:1251–1267.

Kniskern JM, Rausher MD. 2006. Major-gene resistance to the rust pathogen *Coleosporium ipomoeae* is common in natural populations of *Ipomoea purpurea*. New Phytol. 171:137–144.

Kropp BR, Albee S, Flint KM, Zambino P, Szabo L, Thomson S V. 1995. Early detection of systemic rust infections of dyers woad (*Isatis tinctoria*) using the polymerase chain reaction. Weed Sci. 43:467–472.

Kwak IY, Moore CR, Spalding EP, Broman KW. 2014. A simple regression-based method to map quantitative trait loci underlying function-valued phenotypes. Genetics. 197:1409–1416.

Kwak I-Y, Moore CR, Spalding EP, Broman KW. 2016. Mapping quantitative trait loci underlying function-valued traits using functional principal component analysis and multi-trait mapping. G3 Genes, Genomes, Genet. 6:79–86.

Lee DK, Parrish AS, Voigt TB. 2014. Switchgrass and giant *Miscanthus* agronomy. In Engineering and science of biomass feedstock production and provision, pp. 37–59. Springer.

Leimu R, Fischer M. 2008. A meta-analysis of local adaptation in plants. PLoS One. 3:e4010

Lesaffre E, Lawson AB. 2012. Bayesian biostatistics. West Sussex, UK: John Wiley & Sons.

Li F, Upadhyaya NM, Sperschneider J, Matny O, Nguyen-Phuc H, et al., 2019. Emergence of the Ug99 lineage of the wheat stem rust pathogen through somatic hybridisation. Nat. Commun. 10:1–15.

Lovell JT, Shakirov E V, Schwartz S, Lowry DB, Aspinwall MJ, et al., 2016. Promises and challenges of eco-physiological genomics in the field: tests of drought responses in switchgrass. Plant Physiol. 172:734–748.

Lowry DB, Behrman KD, Grabowski P, Morris GP, Kiniry JR, Juenger TE. 2014. Adaptations between ecotypes and along environmental gradients in *Panicum virgatum*. Am. Nat. 183:682–692.

Lowry DB, Lovell JT, Zhang L, Bonnette J, Fay PA, et al., 2019. QTL× environment interactions underlie adaptive divergence in switchgrass across a large latitudinal gradient. Proc. Natl. Acad. Sci. 116:12933–1241.

Mckay JK, Richards JH, Mitchell-Olds T. 2003. Genetics of drought adaptation in *Arabidopsis thaliana*: I. Pleiotropy contributes to genetic correlations among ecological traits. Mol. Ecol. 12:1137–1151.

McNeal FH, Konzak CF, Smith EP, Tate WS, Russell TS. 1971. A uniform system for recording and processing cereal research data. USDA-ARS.

Milano ER, Lowry DB, Juenger TE. 2016. The genetic basis of upland/lowland ecotype divergence in switchgrass (*Panicum virgatum*). G3 Genes, Genomes, Genet. 6:3561–3570.

Morris GP, Grabowski PP, Borevitz JO. 2011. Genomic diversity in switchgrass (*Panicum virgatum*): from the continental scale to a dune landscape. Mol. Ecol. 20:4938–4952.

Mursinoff S, Tack AJM. 2017. Spatial variation in soil biota mediates plant adaptation to a foliar pathogen. New Phytol. 214:644–654.

Parrish DJ, Casler MD, Monti A. 2012. The evolution of switchgrass as an energy crop. Switchgrass, pp. 1–28. Springer.

Peixoto M de M, Sage RF. 2016. Improved experimental protocols to evaluate cold tolerance thresholds in *Miscanthus* and switchgrass rhizomes. GCB Bioenergy. 8:257–268.

Penczykowski RM, Laine A, Koskella B. 2016. Understanding the ecology and evolution of host–parasite interactions across scales. Evol. Appl. 9:37–52.

Price N, Moyers BT, Lopez L, Lasky JR, Monroe JG, et al., 2018. Combining population genomics and fitness QTLs to identify the genetics of local adaptation in *Arabidopsis thaliana*. Proc. Natl. Acad. Sci. 115:5028–5033.

Roelfs AP, Singh RP, Saari EE. 1992. Rust diseases of wheat: concepts and methods of disease management. Mexico, D.F.: CIMMYT. 44–47.

Shaw MW, Osborne TM. 2011. Geographic distribution of plant pathogens in response to climate change. Plant Pathol. 60:31–43.

Simms EL, Triplett J. 1994. Costs and Benefits of Plant Responses to Disease: Resistance and Tolerance. Evolution 48:1973–1985.

Singh A, Knox RE, DePauw RM, Singh AK, Cuthbert RD, et al., 2014. Stripe rust and leaf rust resistance QTL mapping, epistatic interactions, and co-localization with stem rust resistance loci in spring wheat evaluated over three continents. Theor. Appl. Genet. 127:2465–2477.

Singh RP, Hodson DP, Jin Y, Lagudah ES, Ayliffe MA, et al., 2015. Emergence and spread of new races of wheat stem rust fungus: continued threat to food security and prospects of genetic control. Phytopathology. 105:872–884.

Soltani A, MafiMoghaddam S, Oladzad Abbasabadi A, Walter K, Kearns PJ, et al., 2018. Genetic analysis of flooding tolerance in an Andean diversity panel of dry bean (*Phaseolus vulgaris* L.). Front. Plant Sci. 9:1–15.

Thrall PH, Burdon JJ. 2002. Evolution of gene-for-gene systems in metapopulations: the effect of spatial scale of host and pathogen dispersal. Plant Pathol. 51:169–184.

Thrall PH, Laine A, Ravensdale M, Nemri A, Dodds PN, et al., 2012. Rapid genetic change underpins antagonistic coevolution in a natural host-pathogen metapopulation. Ecol. Lett. 15:425–435.

Tian D, Traw MB, Chen JQ, Kreitman M, Bergelson J. 2003. Fitness costs of R-gene-mediated resistance in *Arabidopsis thaliana*. Nature. 423:74–77.

Traw MB, Bergelson J. 2010. Plant immune system incompatibility and the distribution of enemies in natural hybrid zones. Curr. Opin. Plant Biol. 13:466–71.

Uppalapati SR, Serba DD, Ishiga Y, Szabo LJ, Mittal S, et al., 2013. Characterization of the rust fungus, *Puccinia emaculata*, and evaluation of genetic variability for rust resistance in switchgrass populations. Bioenergy Res. 6:458–68.

Vazquez MD, Zemetra R, Peterson CJ, Chen XM, Heesacker A, Mundt CC. 2015. Multi-location wheat stripe rust QTL analysis: genetic background and epistatic interactions. Theor. Appl. Genet. 128:1307–18.

Wickham H. 2016. ggplot2: Elegant Graphics for Data Analysis. Springer-Verlag: New York.

Zeng Q, Wu J, Liu S, Huang S, Wang Q, et al., 2019. A major QTL co-localized on chromosome 6BL and its epistatic interaction for enhanced wheat stripe rust resistance. Theor. Appl. Genet. 132:1409–24.

