## Supplemental Tables and Figures for "Geographic variation in the genetic basis of resistance to leaf rust between locally adapted ecotypes of the biofuel crop switchgrass (*Panicum virgatum*)"

Table S1: Centimorgan positions, maximum SLOD scores, and for significant QTLs. Darker color in the SLOD column indicates strength of QTL.

| Site | Year | Chromosome | Marker | Position (cM) | SLOD | Variation Explained |
|---|---|---|---|---|---|---|
| CLMB | 2016 | 3N | Chr03N_45576768 | 75.78167535 | 2.34 | 6.641523054 |
| CLMB | 2016 | 5K | Chr05K_74431488 | 55.11871179 | 2.58 | 7.348480751 |
| CLMB | 2016 | 9N | Chr09N_83315136 | 72.15246781 | 2.54 | 5.823453642 |
| CLMB | 2017 | 9N | Chr09N_81792576 | 68.94602847 | 3.12 | 4.601569182 |
| CLMB | 2018 | 3N | Chr03N_44605888 | 76.89505827 | 12.06 | 18.2082503 |
| CLMB | 2018 | 9N | Chr09N_89261824 | 78.60470685 | 10.26 | 15.61233461 |
| KBSM | 2016 | 3N | Chr03N_35828928 | 68.69927021 | 2.83 | 7.542549577 |
| KBSM | 2016 | 8K | Chr08K_11288640 | 12.03287596 | 2.21 | 8.857447436 |
| KBSM | 2016 | 9N | Chr09N_89261824 | 78.60470685 | 2.62 | 6.639276268 |
| KBSM | 2017 | 3N | Chr03N_49484736 | 82.10072886 | 4.66 | 7.721438175 |
| KBSM | 2017 | 9N | Chr09N_89261824 | 78.60470685 | 3.64 | 7.360360734 |
| KBSM | 2018 | 1K | Chr01K_64603648 | 63.01696792 | 3.26 | 5.793921956 |
| KBSM | 2018 | 2K | Chr02N_80664320 | 63.69727203 | 3.98 | 5.974580708 |
| KBSM | 2018 | 3K | Chr03K_29874368 | 42.64992868 | 3.69 | 6.150317513 |
| KBSM | 2018 | 3N | Chr03N_34830080 | 67.53941954 | 3.58 | 7.247617966 |
| KBSM | 2018 | 9N | Chr09N_89261824 | 78.60470685 | 7.79 | 17.0689207 |
| KING | 2016 | 1K | Chr01K_50346624 | 42.59984626 | 2.57 | 9.460687778 |
| KING | 2016 | 4K | Chr04K_8737472 | 18.90092347 | 3.76 | 8.593881309 |
| KING | 2016 | 5K | Chr05K_104765312 | 87.31915344 | 2.77 | 8.347244863 |
| KING | 2016 | 7N | Chr07N_55459008 | 33.28497797 | 2.27 | 7.100578559 |
| KING | 2017 | 3N | Chr07N_9934400 | 52.16614119 | 3.08 | 8.659628 |
| KING | 2017 | 4K | Chr04K_8737472 | 18.90092347 | 3.69 | 22.33122934 |
| KING | 2018 | 2K | Chr02K_92526272 | 96.35516997 | 2.99 | 7.684190521 |
| LINC | 2016 | 3N | Chr03N_49103168 | 79.30051735 | 3.61 | 13.32712361 |
| LINC | 2016 | 9K | Chr09K_84580992 | 107.925123 | 3.06 | 6.931193044 |
| LINC | 2016 | 9N | Chr09N_89979776 | 80.67042577 | 4.58 | 12.41654907 |
| MNHT | 2016 | 1K | Chr01K_64043328 | 60.69449872 | 2.36 | 6.255381874 |
| MNHT | 2016 | 3N | Chr03N_45576768 | 75.78167535 | 2.43 | 13.71501936 |
| MNHT | 2016 | 8K | Chr08K_59091712 | 36.03405896 | 2.91 | 12.06194562 |
| MNHT | 2016 | 9K | Chr09K_85136704 | 108.6635311 | 3.63 | 18.6044235 |
| MNHT | 2016 | 9N | Chr09N_91535360 | 84.54263155 | 2.95 | 22.28440992 |
| MNHT | 2017 | 3K | Chr05K_59581632 | 48.33667987 | 4.09 | 8.071250519 |
| MNHT | 2017 | 3N | Chr03N_45187712 | 78.24914389 | 7.37 | 15.29876582 |
| MNHT | 2017 | 8K | Chr08K_11288640 | 12.03287596 | 3.69 | 12.04575939 |
| MNHT | 2017 | 9N | Chr09N_89261824 | 78.60470685 | 4.92 | 10.34664867 |
| OVTN | 2016 | 2K | Chr02K_6118016 | 6.727838082 | 2.19 | 6.496663252 |
| OVTN | 2016 | 9N | Chr09N_13676096 | 26.49110926 | 2.44 | 9.429844595 |
| OVTN | 2017 | 6K | Chr06K_64944768 | 44.50933117 | 3.09 | 8.197845059 |

| | | | | | | |
|---|---|---|---|---|---|---|
| OVTN | 2017 | 8K | Chr08K_22265600 | 19.52389775 | 2.83 | 7.181421805 |
| OVTN | 2018 | 2N | Chr02N_42430400 | 42.12622142 | 3.37 | 7.141948354 |
| PKLE | 2018 | 1K | Chr01K_62280768 | 58.12575861 | 3.93 | 8.014268212 |
| PKLE | 2018 | 2N | Chr02N_87675008 | 74.10375844 | 3.32 | 6.527790772 |
| TMPL | 2016 | 1K | Chr01K_50346624 | 42.59984626 | 2.18 | 9.524698032 |
| TMPL | 2016 | 3K | Chr03K_28313344 | 40.93920108 | 2.67 | 7.345370404 |
| TMPL | 2016 | 5N | Chr05N_94241280 | 78.45331178 | 2.34 | 9.93342625 |
| TMPL | 2016 | 9K | Chr09K_85136704 | 108.6635311 | 2.12 | 9.584801905 |
| TMPL | 2017 | 3K | Chr01K_38848832 | 47.3236671 | 4.07 | 8.721804146 |
| TMPL | 2017 | 3N | Chr03N_35828928 | 68.69927021 | 7.21 | 24.23242194 |
| TMPL | 2017 | 8N | Chr08N_22149056 | 22.0240102 | 3.31 | 6.564604856 |
| TMPL | 2017 | 9N | Chr09N_88105088 | 75.30296451 | 3.19 | 10.29637002 |
| TMPL | 2018 | 2N | Chr02N_57893888 | 47.8728519 | 4.39 | 14.14141462 |

Table S2: ANOVA tests for epistatic interactions between 3N and 9N QTLs. Bold values are significant at Bonferroni-corrected p-value of 0.00263.

| DF | SS | MS | F | P | SITE | Factor |
|---|---|---|---|---|---|---|
| 1 | 7.183 | 7.183 | 2.599 | 0.1079 | CLMB_2016 | 3N |
| **1** | **35.501** | **35.501** | **12.844** | **3.88E-04** | **CLMB_2016** | **9N** |
| 1 | 0.794 | 0.794 | 0.287 | 0.5923 | CLMB_2016 | Interaction |
| 341 | 942.511 | 2.764 | | | CLMB_2016 | Residuals |
| 1 | 0.658 | 0.658 | 0.415 | 0.5200 | CLMB_2017 | 3N |
| 1 | 10.874 | 10.874 | 6.852 | 0.0092 | CLMB_2017 | 9N |
| 1 | 2.633 | 2.633 | 1.659 | 0.1986 | CLMB_2017 | Interaction |
| 345 | 547.531 | 1.587 | | | CLMB_2017 | Residuals |
| **1** | **140.758** | **140.758** | **59.046** | **1.60E-13** | **CLMB_2018** | **3N** |
| **1** | **86.360** | **86.360** | **36.227** | **4.48E-09** | **CLMB_2018** | **9N** |
| **1** | **70.828** | **70.828** | **29.711** | **9.55E-08** | **CLMB_2018** | **Interaction** |
| 346 | 824.817 | 2.384 | | | CLMB_2018 | Residuals |
| 1 | 0.150 | 0.150 | 0.065 | 0.7992 | KBSM_2016 | 3N |
| 1 | 7.933 | 7.933 | 3.439 | 0.0645 | KBSM_2016 | 9N |
| 1 | 0.633 | 0.633 | 0.274 | 0.6008 | KBSM_2016 | Interaction |
| 346 | 798.213 | 2.307 | | | KBSM_2016 | Residuals |
| 1 | 4.114 | 4.114 | 3.463 | 0.0636 | KBSM_2017 | 3N |
| **1** | **19.647** | **19.647** | **16.539** | **5.92E-05** | **KBSM_2017** | **9N** |
| 1 | 0.574 | 0.574 | 0.483 | 0.4875 | KBSM_2017 | Interaction |
| 341 | 405.097 | 1.188 | | | KBSM_2017 | Residuals |
| **1** | **94.779** | **94.779** | **13.215** | **3.20E-04** | **KBSM_2018** | **3N** |
| **1** | **348.128** | **348.128** | **48.541** | **1.68E-11** | **KBSM_2018** | **9N** |
| **1** | **72.738** | **72.738** | **10.142** | **0.0016** | **KBSM_2018** | **Interaction** |
| 341 | 2445.613 | 7.172 | | | KBSM_2018 | Residuals |
| 1 | 0.033 | 0.033 | 0.011 | 0.9150 | KING_2016 | 3N |
| 1 | 0.001 | 0.001 | 0.000 | 0.9859 | KING_2016 | 9N |
| 1 | 8.743 | 8.743 | 3.008 | 0.0839 | KING_2016 | Interaction |
| 304 | 883.674 | 2.907 | | | KING_2016 | Residuals |
| 1 | 1.068 | 1.068 | 0.441 | 0.5070 | KING_2017 | 3N |
| 1 | 3.550 | 3.550 | 1.467 | 0.2268 | KING_2017 | 9N |
| 1 | 1.944 | 1.944 | 0.803 | 0.3709 | KING_2017 | Interaction |
| 299 | 723.813 | 2.421 | | | KING_2017 | Residuals |
| 1 | 2.012 | 2.012 | 0.306 | 0.5803 | KING_2018 | 3N |
| 1 | 8.376 | 8.376 | 1.276 | 0.2597 | KING_2018 | 9N |
| 1 | 0.576 | 0.576 | 0.088 | 0.7672 | KING_2018 | Interaction |
| 256 | 1680.176 | 6.563 | | | KING_2018 | Residuals |
| **1** | **46.963** | **46.963** | **14.639** | **1.59E-04** | **LINC_2016** | **3N** |
| **1** | **85.309** | **85.309** | **26.592** | **4.66E-07** | **LINC_2016** | **9N** |

| | | | | | | |
|---|---|---|---|---|---|---|
| **1** | **32.653** | **32.653** | **10.178** | **0.0016** | **LINC_2016** | **Interaction** |
| 290 | 930.356 | 3.208 | | | LINC_2016 | Residuals |
| 1 | 11.236 | 11.236 | 3.956 | 0.0476 | MNHT_2016 | 3N |
| **1** | **53.284** | **53.284** | **18.760** | **2.01E-05** | **MNHT_2016** | **9N** |
| 1 | 1.650 | 1.650 | 0.581 | 0.4466 | MNHT_2016 | Interaction |
| 308 | 874.802 | 2.840 | | | MNHT_2016 | Residuals |
| **1** | **9.381** | **9.381** | **21.846** | **4.41E-06** | **MNHT_2017** | **3N** |
| **1** | **7.544** | **7.544** | **17.568** | **3.62E-05** | **MNHT_2017** | **9N** |
| **1** | **5.784** | **5.784** | **13.471** | **2.85E-04** | **MNHT_2017** | **Interaction** |
| 311 | 133.545 | 0.429 | | | MNHT_2017 | Residuals |
| 1 | 5.835 | 5.835 | 5.409 | 0.0208 | OVTN_2016 | 3N |
| 1 | 4.058 | 4.058 | 3.762 | 0.0535 | OVTN_2016 | 9N |
| 1 | 1.253 | 1.253 | 1.161 | 0.2821 | OVTN_2016 | Interaction |
| 274 | 295.574 | 1.079 | | | OVTN_2016 | Residuals |
| 1 | 5.227 | 5.227 | 2.742 | 0.0988 | OVTN_2017 | 3N |
| 1 | 11.383 | 11.383 | 5.971 | 0.0151 | OVTN_2017 | 9N |
| 1 | 0.665 | 0.665 | 0.349 | 0.5552 | OVTN_2017 | Interaction |
| 305 | 581.431 | 1.906 | | | OVTN_2017 | Residuals |
| 1 | 2.517 | 2.517 | 1.521 | 0.2190 | PKLE_2016 | 3N |
| 1 | 1.284 | 1.284 | 0.776 | 0.3796 | PKLE_2016 | 9N |
| 1 | 3.358 | 3.358 | 2.030 | 0.1560 | PKLE_2016 | Interaction |
| 179 | 296.099 | 1.654 | | | PKLE_2016 | Residuals |
| 1 | 1.195 | 1.195 | 0.365 | 0.5464 | PKLE_2018 | 3N |
| 1 | 0.649 | 0.649 | 0.198 | 0.6567 | PKLE_2018 | 9N |
| 1 | 0.478 | 0.478 | 0.146 | 0.7027 | PKLE_2018 | Interaction |
| 336 | 1101.400 | 3.278 | | | PKLE_2018 | Residuals |
| 1 | 0.716 | 0.716 | 1.866 | 0.1730 | TMPL_2016 | 3N |
| 1 | 0.875 | 0.875 | 2.278 | 0.1322 | TMPL_2016 | 9N |
| 1 | 1.117 | 1.117 | 2.910 | 0.0891 | TMPL_2016 | Interaction |
| 301 | 115.542 | 0.384 | | | TMPL_2016 | Residuals |
| **1** | **274.980** | **274.980** | **62.623** | **5.18E-14** | **TMPL_2017** | **3N** |
| **1** | **98.956** | **98.956** | **22.536** | **3.23E-06** | **TMPL_2017** | **9N** |
| **1** | **124.991** | **124.991** | **28.465** | **1.91E-07** | **TMPL_2017** | **Interaction** |
| 293 | 1286.574 | 4.391 | | | TMPL_2017 | Residuals |
| 1 | 0.003 | 0.003 | 0.001 | 0.9821 | TMPL_2018 | 3N |
| 1 | 50.785 | 50.785 | 7.935 | 0.0052 | TMPL_2018 | 9N |
| 1 | 24.132 | 24.132 | 3.770 | 0.0531 | TMPL_2018 | Interaction |
| 305 | 1952.090 | 6.400 | | | TMPL_2018 | Residuals |

Table S3: Genes in 10% LOD-drop interval for *Prr1* and *Prr2*. Links to each gene in the JGI Phytozome can be found by pasting the value in the Link Suffix column to:
"https://phytozome-next.jgi.doe.gov/report/transcript/Pvirgatum_v5_1/Pavir. "

| Name | Position | Link Suffix | Annotation |
|---|---|---|---|
| Pavir.3NG156394 | 45021487 | 3NG156394.1 | NA |
| Pavir.3NG156300 | 45009687 | 3NG156300.1 | NA |
| Pavir.3NG160194 | 44899275 | 3NG160194.1 | SWR1-complex protein 5/Craniofacial development protein |
| Pavir.3NG160100 | 44743166 | 3NG160100.1 | Pinoresinol/lariciresinol reductase |
| Pavir.3NG160000 | 44738363 | 3NG160000.1 | Pinoresinol/lariciresinol reductase |
| Pavir.3NG168300 | 44704748 | 3NG168300.2 | PROTEIN S-ACYLTRANSFERASE 18 |
| Pavir.3NG168400 | 44693194 | 3NG168400.1 | Glutathione synthetase |
| Pavir.3NG168600 | 44641957 | 3NG168600.1 | NA |
| Pavir.3NG168900 | 44629466 | 3NG168900.2 | MULTI-COPPER OXIDASE |
| Pavir.3NG169000 | 44590591 | 3NG169000.1 | F26K24.10 PROTEIN |
| Pavir.3NG168576 | 44564470 | 3NG168576.1 | NA |
| Pavir.9NG482800 | 54120780 | 9NG482800.1 | Hydrophobic seed protein (Hydrophob_seed) |
| Pavir.9NG482700 | 54113485 | 9NG482700.1 | Hydrophobic seed protein (Hydrophob_seed) |
| Pavir.9NG482600 | 54109974 | 9NG482600.2 | small subunit ribosomal protein S15 (RP-S15, MRPS15, rpsO) |
| Pavir.9NG482500 | 54107577 | 9NG482500.3 | 3-HYDROXYISOBUTYRYL-COA HYDROLASE-LIKE PROTEIN 1, MITOCHONDRIAL |
| Pavir.9NG482400 | 54102571 | 9NG482400.1 | TREHALOSE-PHOSPHATE PHOSPHATASE F-RELATED |
| Pavir.9NG482300 | 54097773 | 9NG482300.1 | NA |
| Pavir.9NG482100 | 54049477 | 9NG482100.2 | large subunit ribosomal protein L44e (RP-L44e, RPL44) |
| Pavir.9NG482000 | 54048146 | 9NG482000.1 | NA |
| Pavir.9NG481800 | 54039738 | 9NG481800.1 | NA |
| Pavir.9NG481700 | 54035257 | 9NG481700.1 | EXOSTOSIN HEPARAN SULFATE GLYCOSYLTRANSFERASE -RELATED |
| Pavir.9NG481600 | 54031703 | 9NG481600.1 | FLAVIN-CONTAINING MONOOXYGENASE FMO GS-OX-LIKE 1-RELATED |
| Pavir.9NG481500 | 54029233 | 9NG481500.2 | NA |
| Pavir.9NG481400 | 54013936 | 9NG481400.1 | Plant protein of unknown function (DUF863) (DUF863) |
| Pavir.9NG481300 | 54003804 | 9NG481300.1 | Poly(ADP-ribose) polymerase, catalytic domain |
| Pavir.9NG481200 | 53985268 | 9NG481200.1 | OXYSTEROL-BINDING PROTEIN-RELATED |
| Pavir.9NG481100 | 53929119 | 9NG481100.1 | OLIGOPEPTIDE TRANSPORTER-RELATED |
| Pavir.9NG481000 | 53900565 | 9NG481000.1 | OLIGOPEPTIDE TRANSPORTER-RELATED |
| Pavir.9NG480900 | 53888536 | 9NG480900.2 | DIMETHYLANILINE MONOOXYGENASE |
| Pavir.9NG480800 | 53886187 | 9NG480800.1 | DNA (cytosine-5-)-methyltransferase / Type II DNA methylase |
| Pavir.9NG480700 | 53880616 | 9NG480700.1 | Protease inhibitor/seed storage/LTP family (Tryp_alpha_amyl) |
| Pavir.9NG480500 | 53854808 | 9NG480500.2 | CYSTEINE-TYPE PEPTIDASE |
| Pavir.9NG477528 | 53851136 | 9NG477528.1 | NA |
| Pavir.9NG477514 | 53844787 | 9NG477514.1 | DIMETHYLANILINE MONOOXYGENASE |
| Pavir.9NG477500 | 53732633 | 9NG477500.1 | DIMETHYLANILINE MONOOXYGENASE |
| Pavir.9NG477400 | 53729995 | 9NG477400.3 | FAMILY NOT NAMED // QWRF MOTIF-CONTAINING PROTEIN 7 |
| Pavir.9NG477100 | 53646456 | 9NG477100.1 | GLYCOGENIN |
| Pavir.9NG477000 | 53640997 | 9NG477000.5 | SF2 - HOLLIDAY JUNCTION RESOLVASE-RELATED |
| Pavir.9NG476800 | 53633004 | 9NG476800.1 | C2H2-LIKE ZINC FINGER PROTEIN-RELATED |

| | | | |
|---|---|---|---|
| Pavir.9NG476600 | 53597763 | 9NG476600.1 | Rare lipoprotein A (RlpA)-like double-psi beta-barrel (DPBB_1) |
| Pavir.9NG476228 | 53579740 | 9NG476228.1 | NA |
| Pavir.9NG476214 | 53576135 | 9NG476214.1 | NA |
| Pavir.9NG476200 | 53572901 | 9NG476200.1 | Rare lipoprotein A (RlpA)-like double-psi beta-barrel (DPBB_1) |
| Pavir.9NG476100 | 53564120 | 9NG476100.1 | Rare lipoprotein A (RlpA)-like double-psi beta-barrel (DPBB_1) |
| Pavir.9NG476000 | 53552178 | 9NG476000.1 | golgi-specific brefeldin A-resistance guanine nucleotide exchange factor 1 (GBF1) |
| Pavir.9NG475900 | 53528529 | 9NG475900.1 | Rare lipoprotein A (RlpA)-like double-psi beta-barrel (DPBB_1) |
| Pavir.9NG475800 | 53498024 | 9NG475800.1 | Rare lipoprotein A (RlpA)-like double-psi beta-barrel (DPBB_1) |
| Pavir.9NG475542 | 53491219 | 9NG475542.1 | Immunoglobulin-like fold |
| Pavir.9NG475528 | 53484325 | 9NG475528.1 | Pollen allergen (Pollen_allerg_1) |
| Pavir.9NG475514 | 53482114 | 9NG475514.1 | Rare lipoprotein A (RlpA)-like double-psi beta-barrel (DPBB_1) |
| Pavir.9NG475500 | 53468411 | 9NG475500.1 | Rare lipoprotein A (RlpA)-like double-psi beta-barrel (DPBB_1) |
| Pavir.9NG475400 | 53454913 | 9NG475400.1 | SENTRIN/SUMO-SPECIFIC PROTEASE // GH15225P |
| Pavir.9NG475214 | 53449579 | 9NG475214.1 | NA |
| Pavir.9NG475200 | 53445456 | 9NG475200.1 | Rare lipoprotein A (RlpA)-like double-psi beta-barrel (DPBB_1) |
| Pavir.9NG474700 | 53422764 | 9NG474700.1 | Rare lipoprotein A (RlpA)-like double-psi beta-barrel (DPBB_1) |
| Pavir.9NG474600 | 53416180 | 9NG474600.1 | STEROL REGULATORY ELEMENT-BINDING PROTEIN |
| Pavir.9NG474500 | 53381755 | 9NG474500.2 | CCR4-NOT transcription complex subunit 1 (CNOT1, NOT1) |
| Pavir.9NG483114 | 53328029 | 9NG483114.1 | NA |
| Pavir.9NG483100 | 53325843 | 9NG483100.1 | Domain of unknown function (DUF239) |

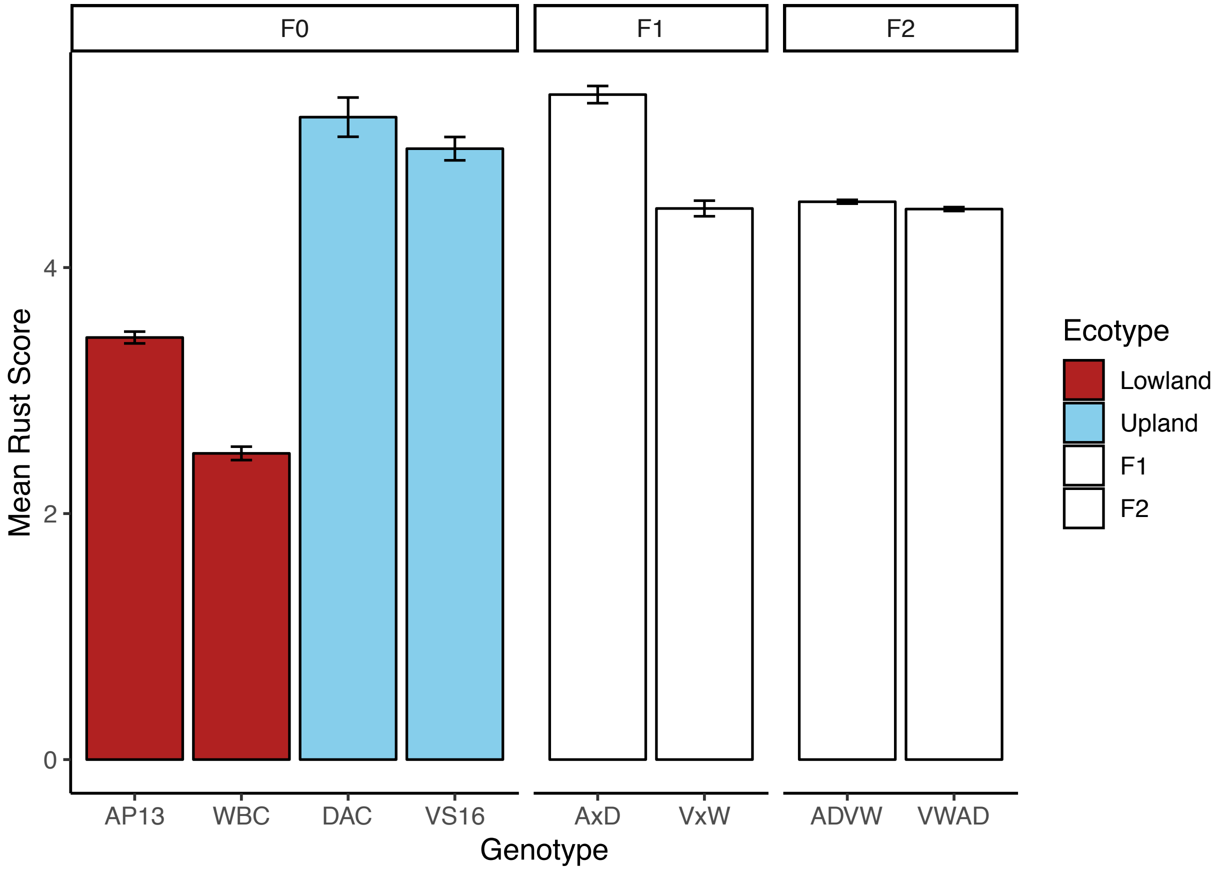

**Figure S1**: Mean rust scores in different generations of the mapping cross over all sites and years. The F0 generation includes the grandparental lines from the two ecotypes: AP13, WBC, VS16, and DAC. The F1 generation includes the two crosses, AxD (AP13 x DAC) and VxW (VS16 x WBC). The F2 generation includes the mapping population, with cytoplasm from either the upland lineage (ADVW) or the lowland lineage (VWAD). Error bars show SE.

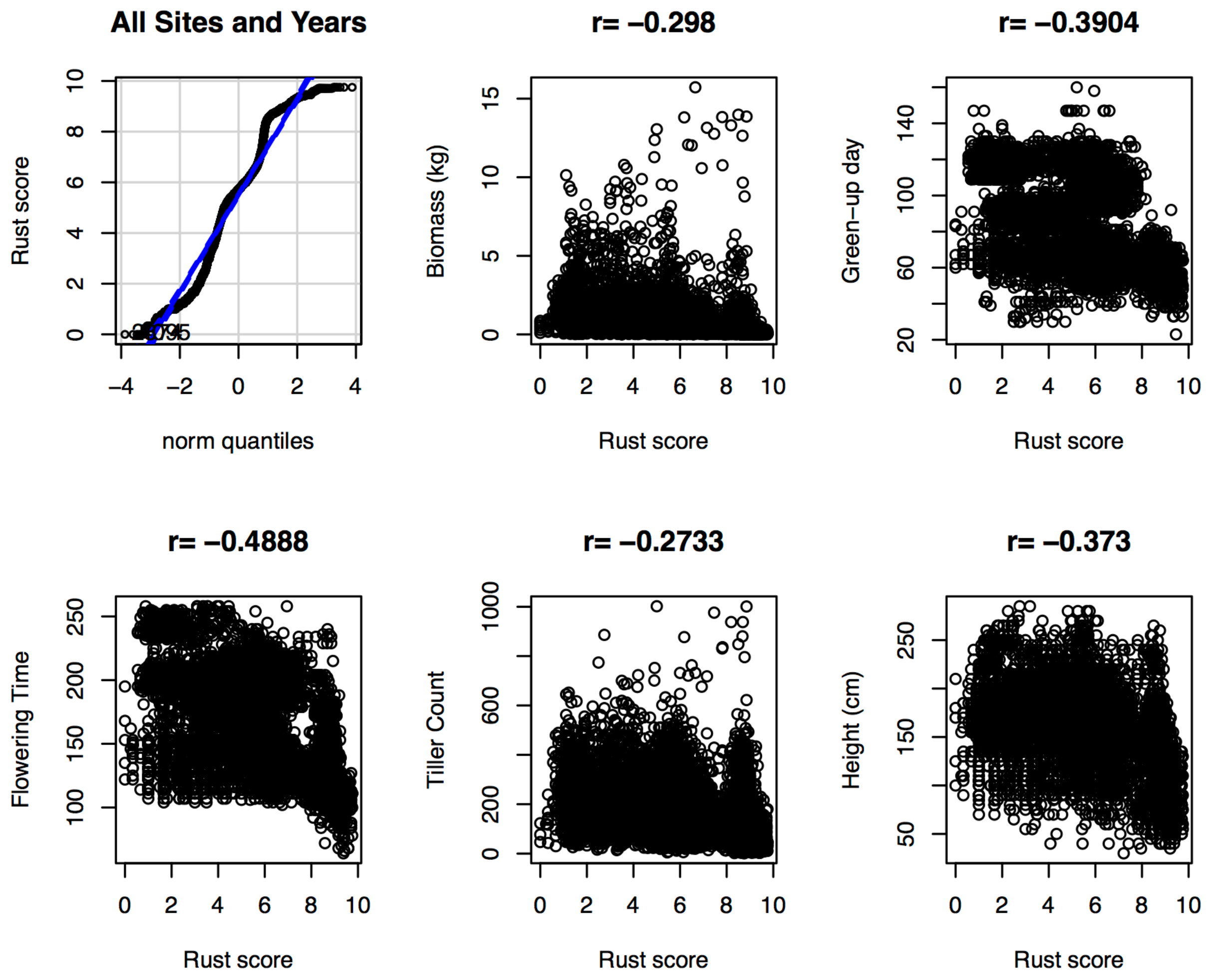

**Figure S2**: Correlations between mean rust score and several morphological and phenology traits across all sites and years. The upper left panel shows the quantile-quantile plot for rust scores. Pearson's r values are shown at the top of each plot. $P \ll 0.0001$ for all correlation tests.

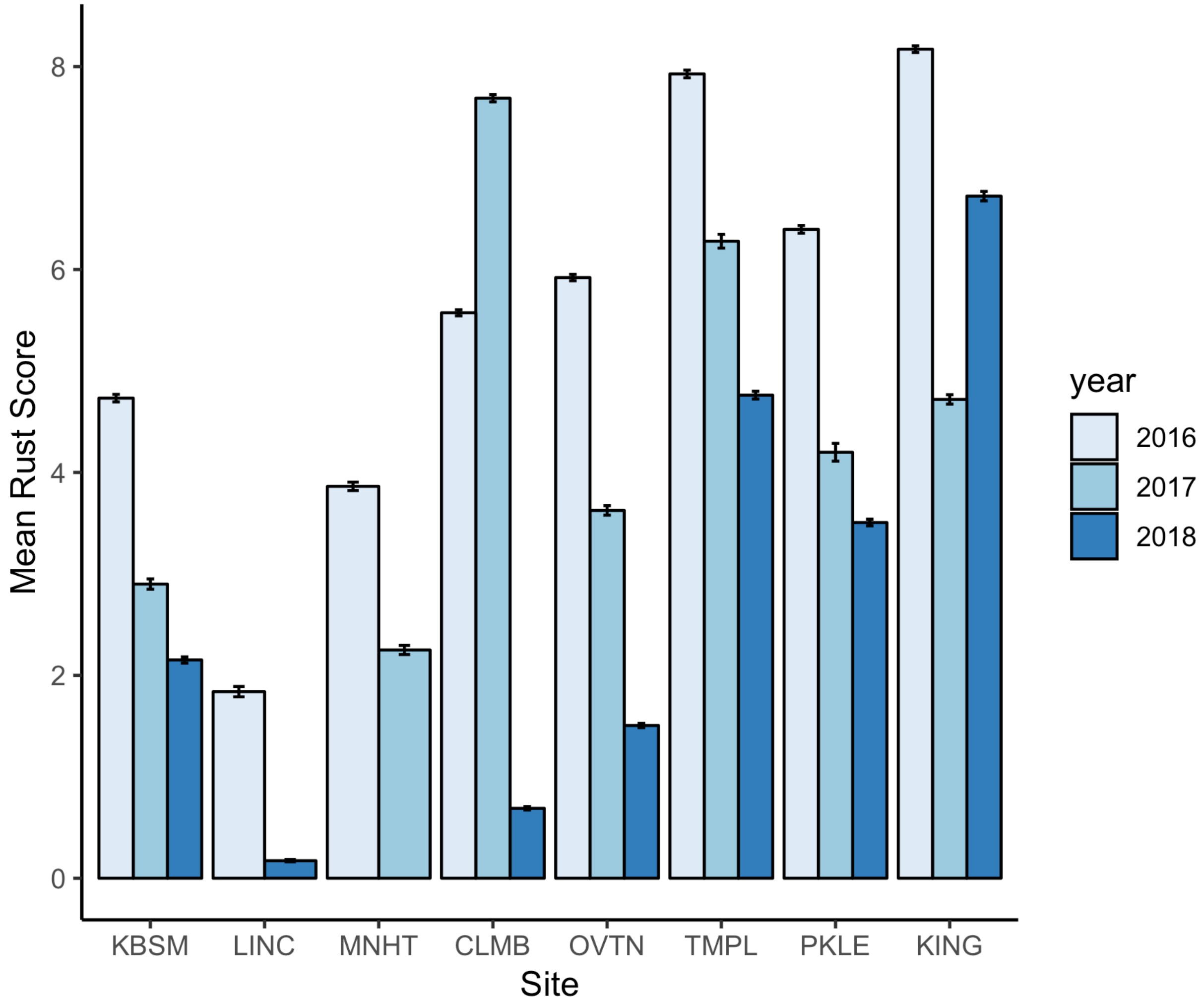

**Figure S3**: Mean rust score subset by site and year showing a decline in most sites over time. Sites are ordered by latitude, from northern KBSM to southern KING. Data for LINC in 2017 and MNHT in 2018 were not collected due to technical issues. Error bars show SE.

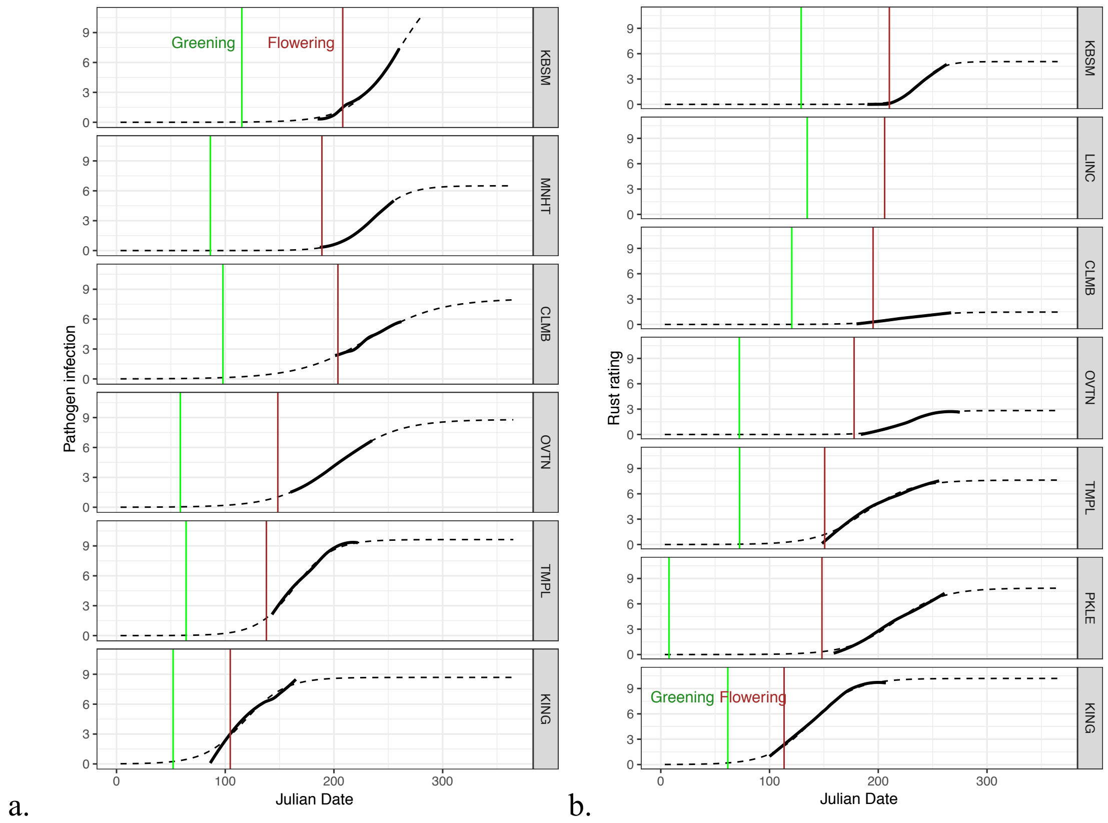

**Figure S4**: Rust progression curves in a. 2017 and b. 2018. Black lines show smoothed mean rust values for sampled dates, black dotted lines show fitted logistic curves to sampled data. Green and brown vertical lines show green-up date and date of first flowering, respectively.

**Figure S5**: Time-Series data for additional sites and years

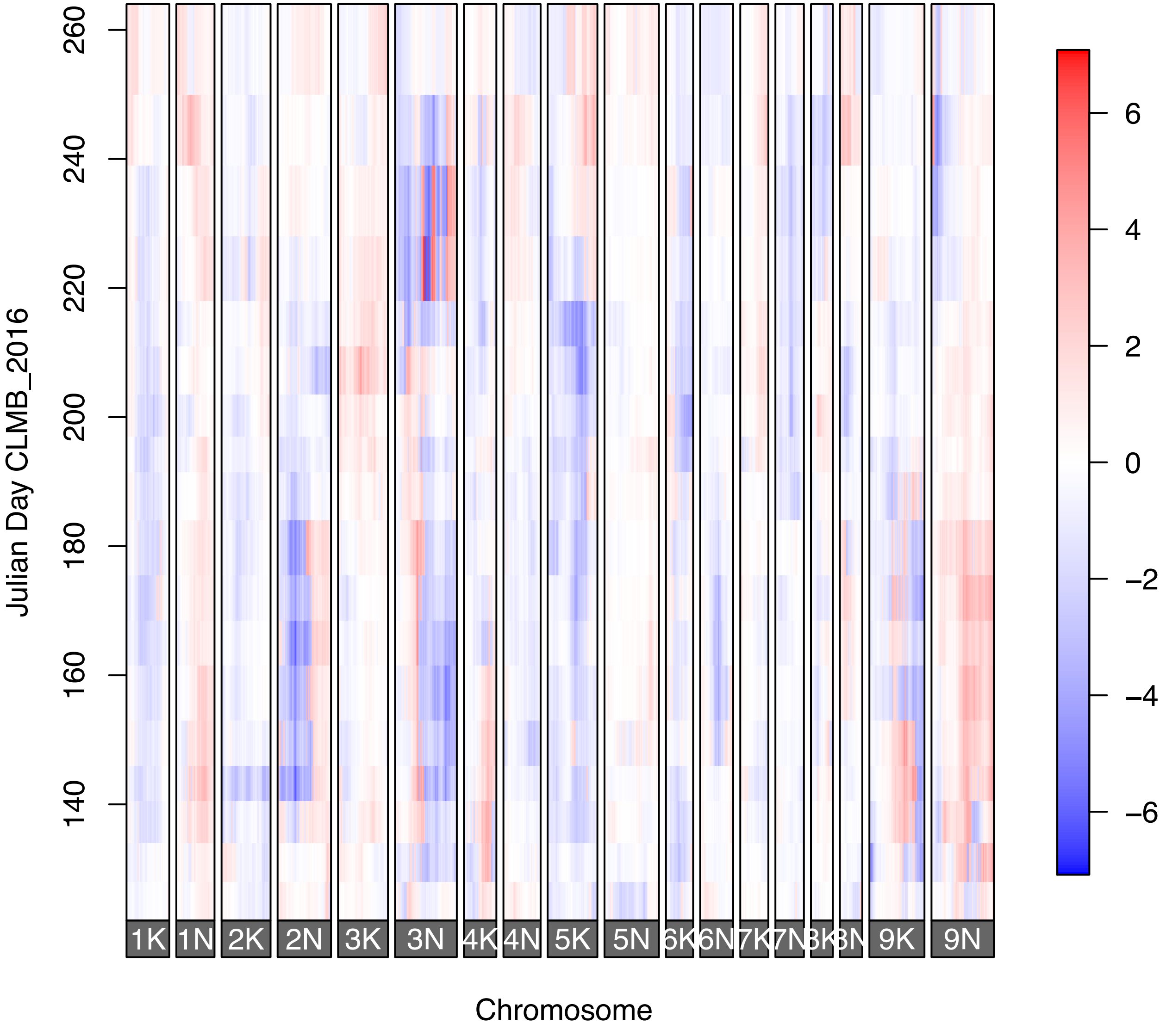

Julian Day CLMB_2016
260
240
220
200
180
160
140
1K
1N
2K
2N
3K
3N
4K
4N
5K
5N
6N
9K
9N
Chromosome
6
4
2
0
−2
−4
−6

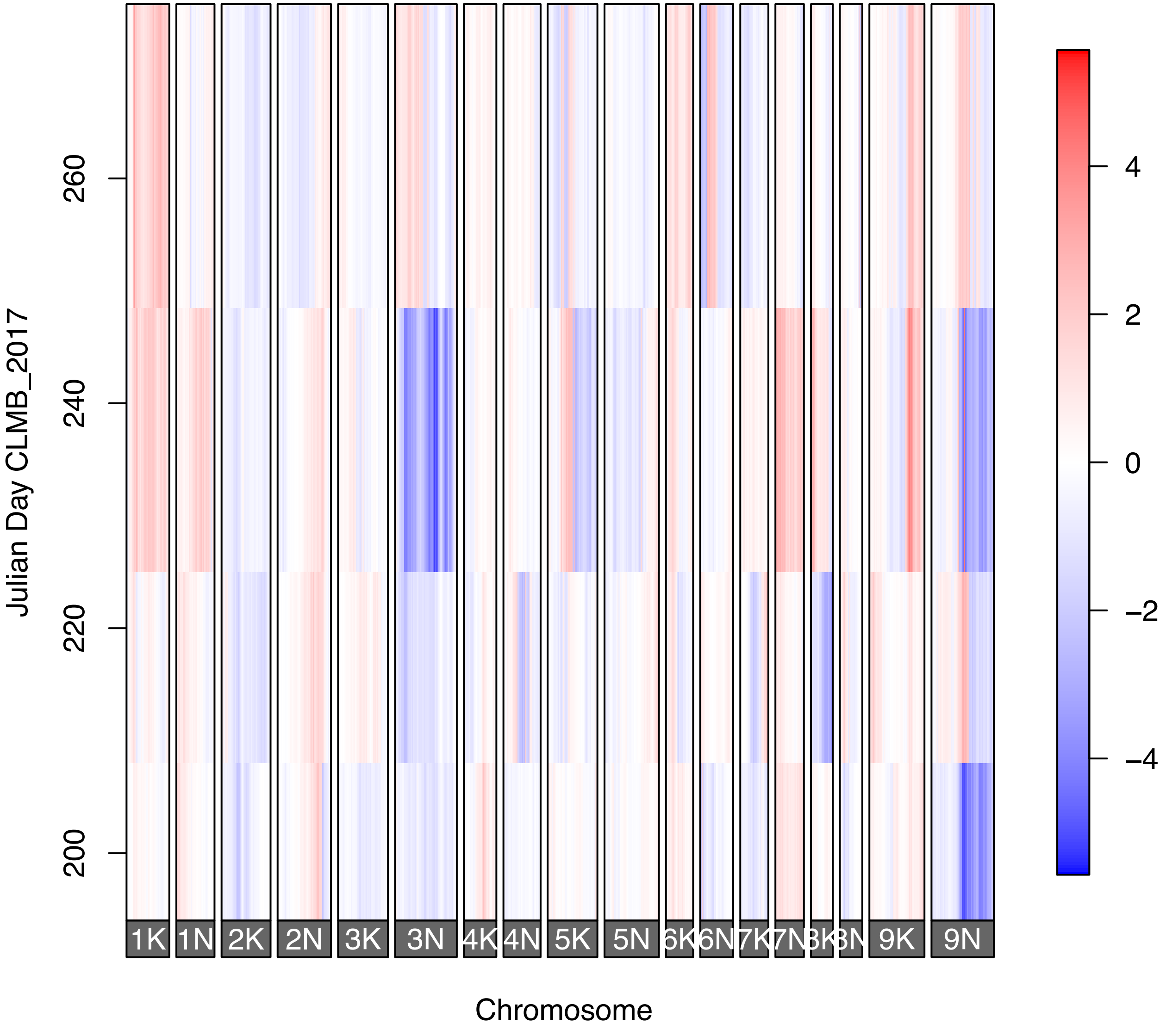

Julian Day CLMB_2017
200
220
240
260
1K
1N
2K
2N
3K
3N
4K
4N
5K
5N
6K
6N
7K
7N
9K
9N
Chromosome
4
2
0
−2
−4

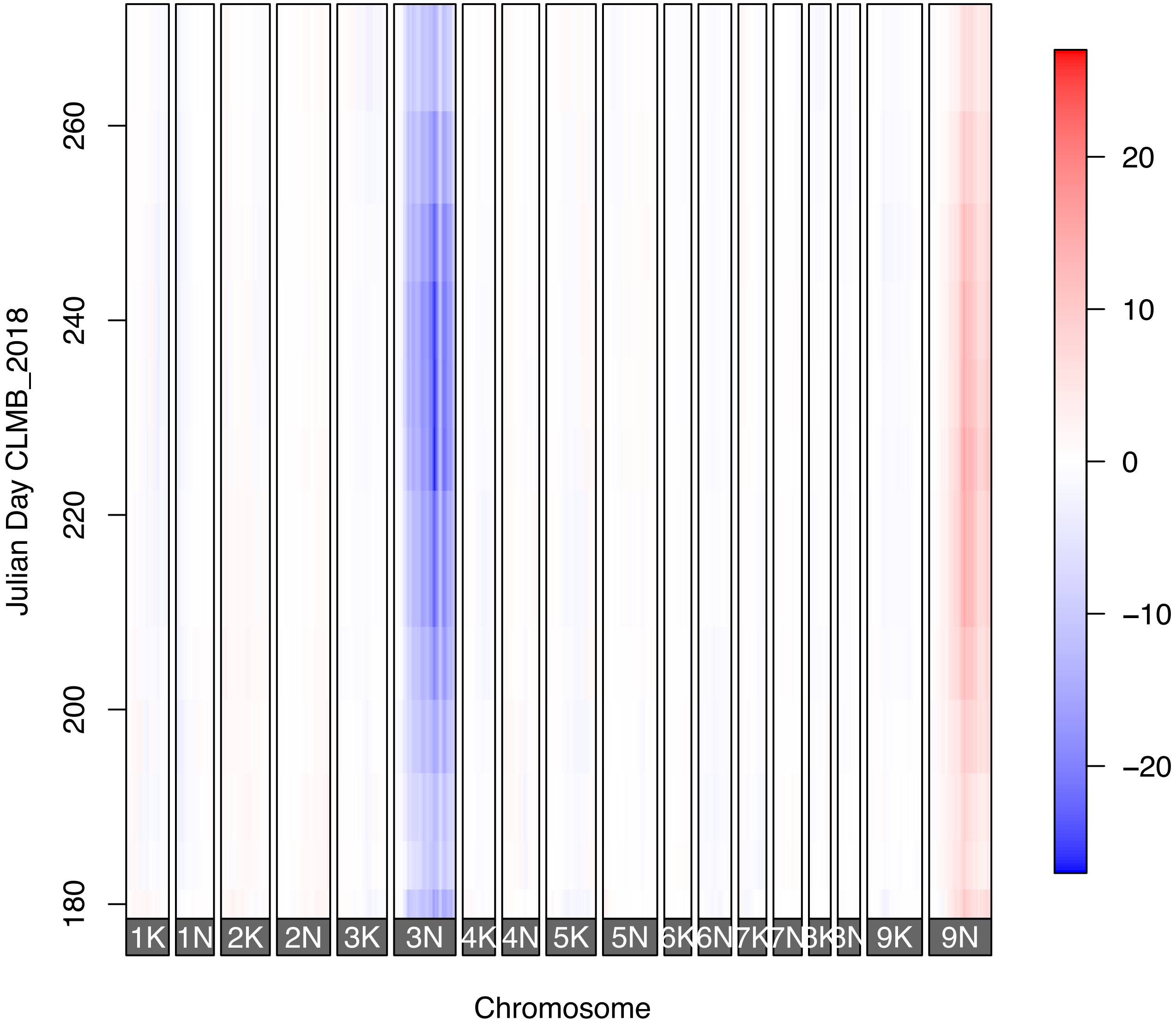
Julian Day CLMB_2018
180
200
220
240
260
1K
1N
2K
2N
3K
3N
4K
4N
5K
5N
6K
6N
7K
7N
8K
8N
9K
9N
Chromosome
20
10
0
−10
−20

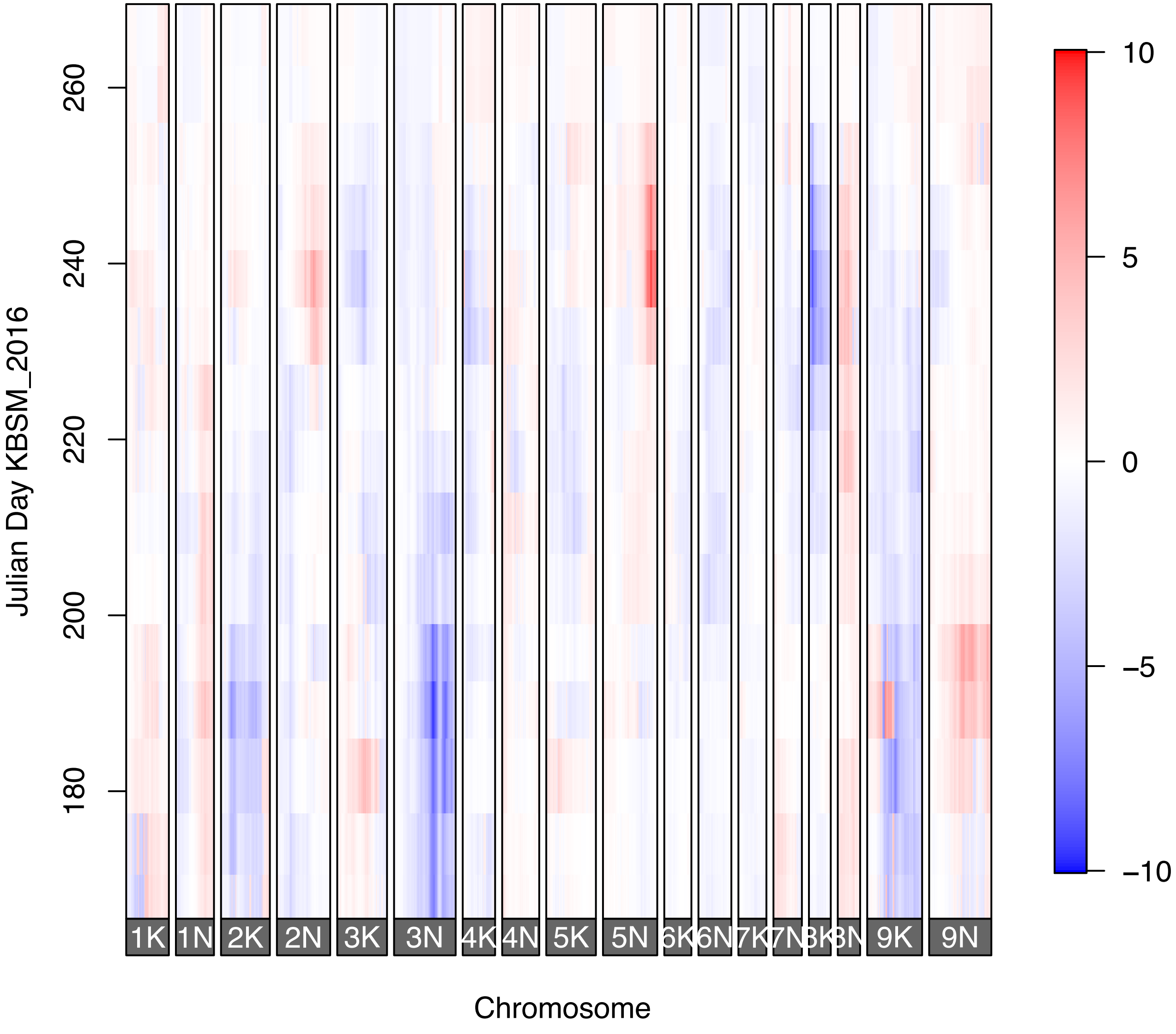

Julian Day KBSM_2016
180
200
220
240
260
1K
1N
2K
2N
3K
3N
4K
4N
5K
5N
6K
6N
7K
7N
8K
8N
9K
9N
Chromosome
10
5
0
−5
−10

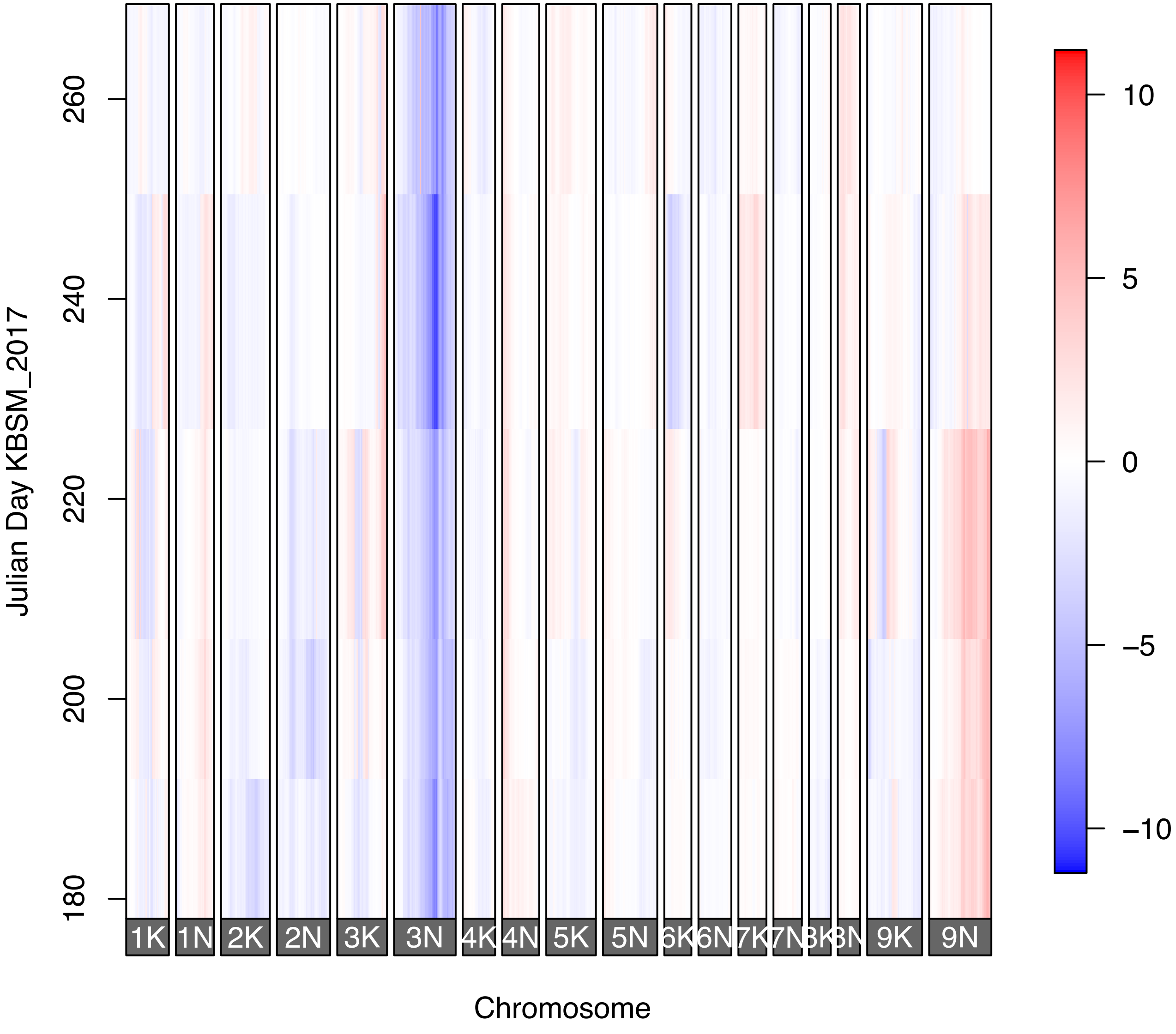
Julian Day KBSM_2017
180
200
220
240
260
1K
1N
2K
2N
3K
3N
4K
4N
5K
5N
6K
6N
7K
7N
9K
9N
Chromosome
10
5
0
−5
−10

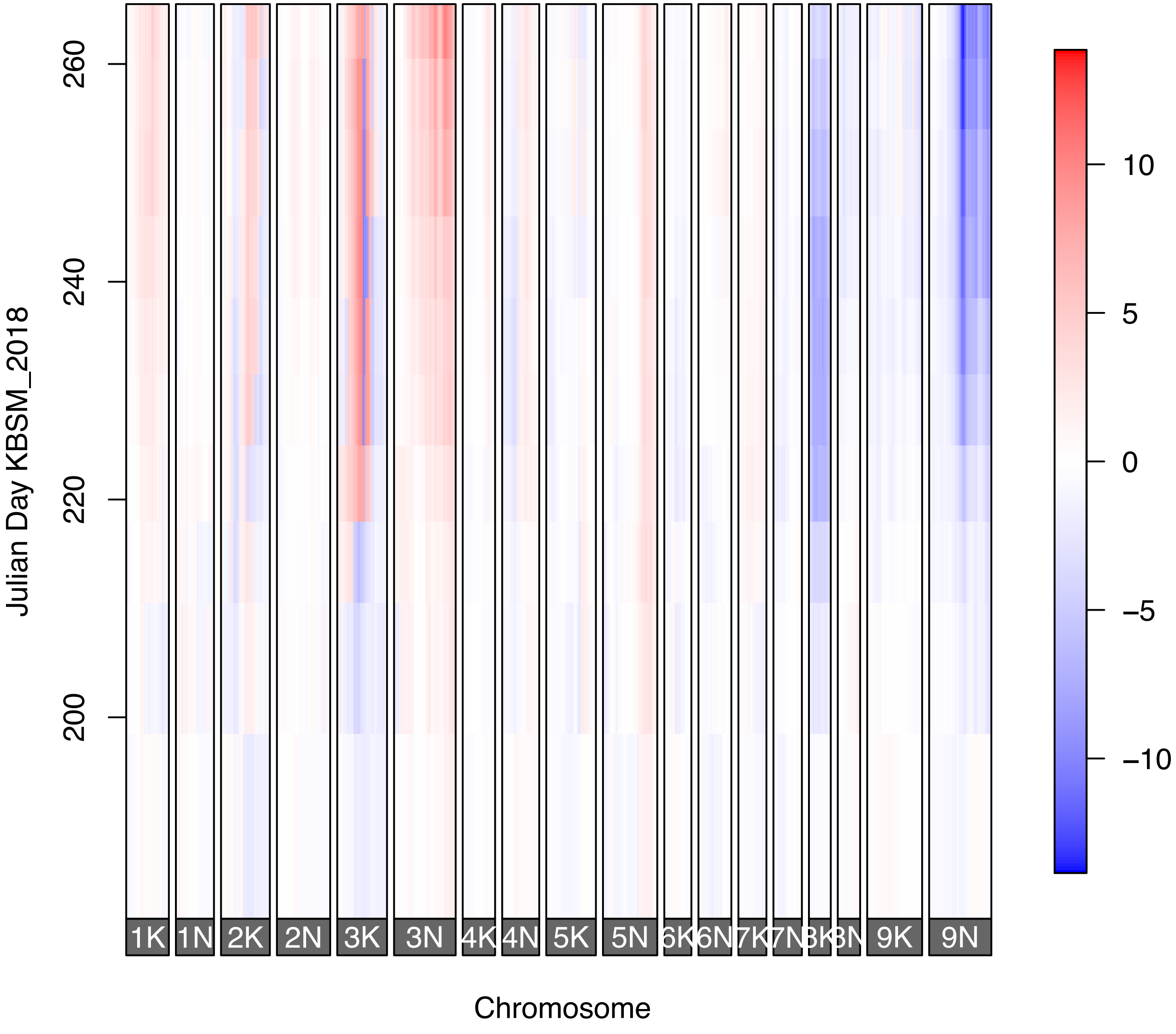

Julian Day KBSM_2018
260
240
220
200
1K
1N
2K
2N
3K
3N
4K
4N
5K
5N
6K
6N
7K
7N
8K
8N
9K
9N
Chromosome
10
5
0
−5
−10

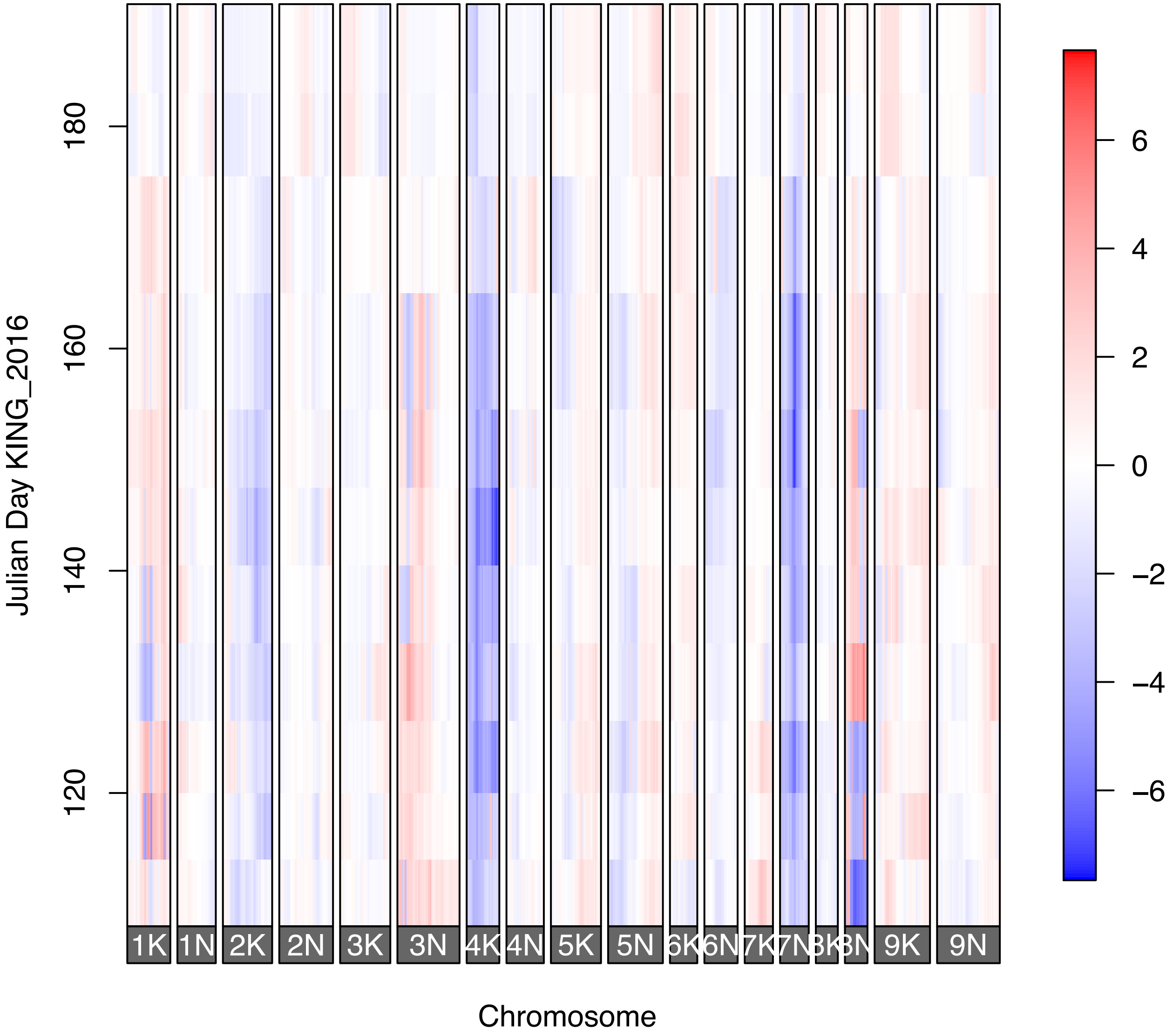
Julian Day KING_2016
180
160
140
120
1K
1N
2K
2N
3K
3N
4K
4N
5K
5N
6K
6N
7K
7N
8K
8N
9K
9N
Chromosome
6
4
2
0
−2
−4
−6

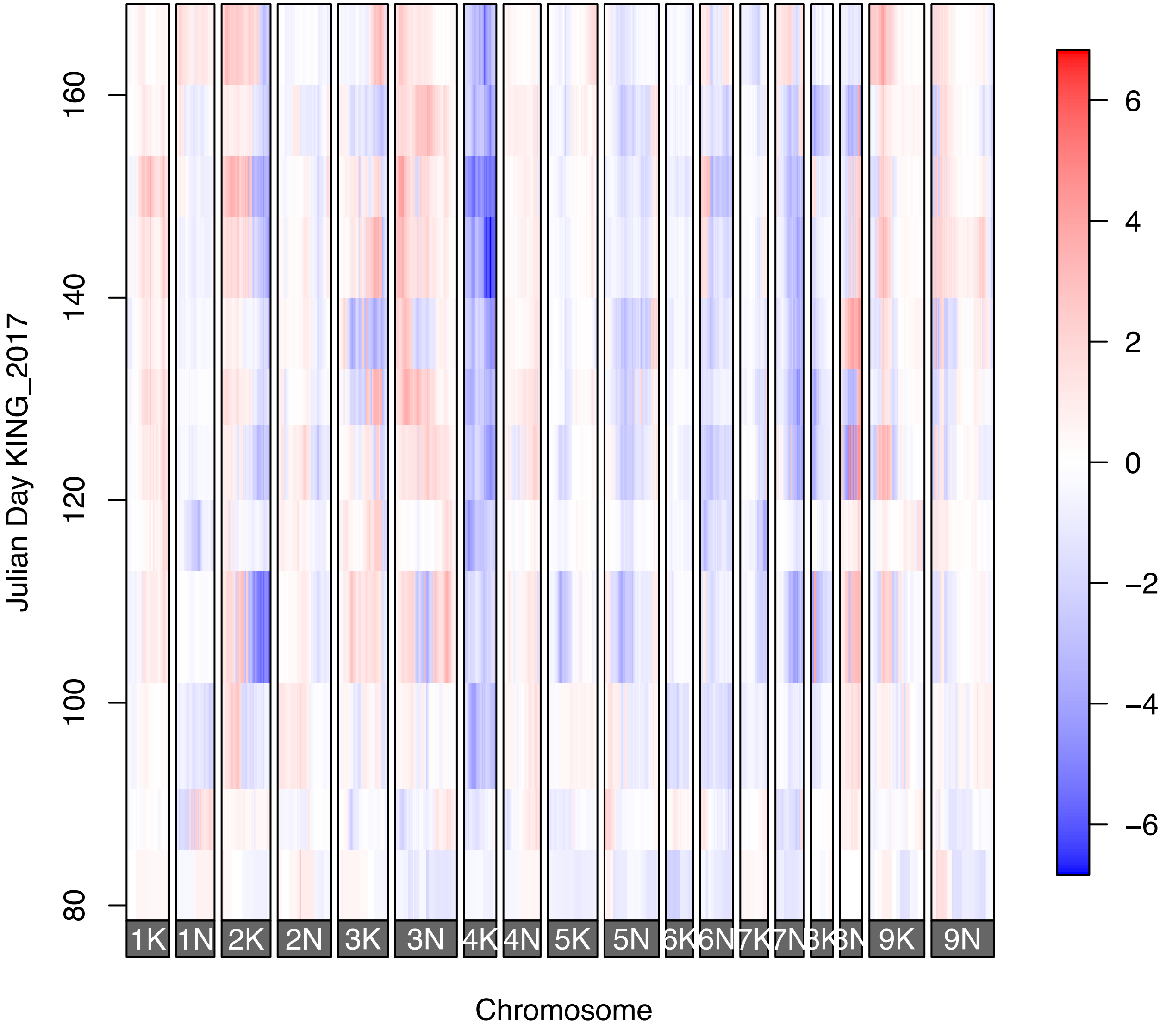

Julian Day KING_2017
80
100
120
140
160
1K
1N
2K
2N
3K
3N
4K
4N
5K
5N
6K
6N
7K
7N
8K
8N
9K
9N
Chromosome
6
4
2
0
−2
−4
−6

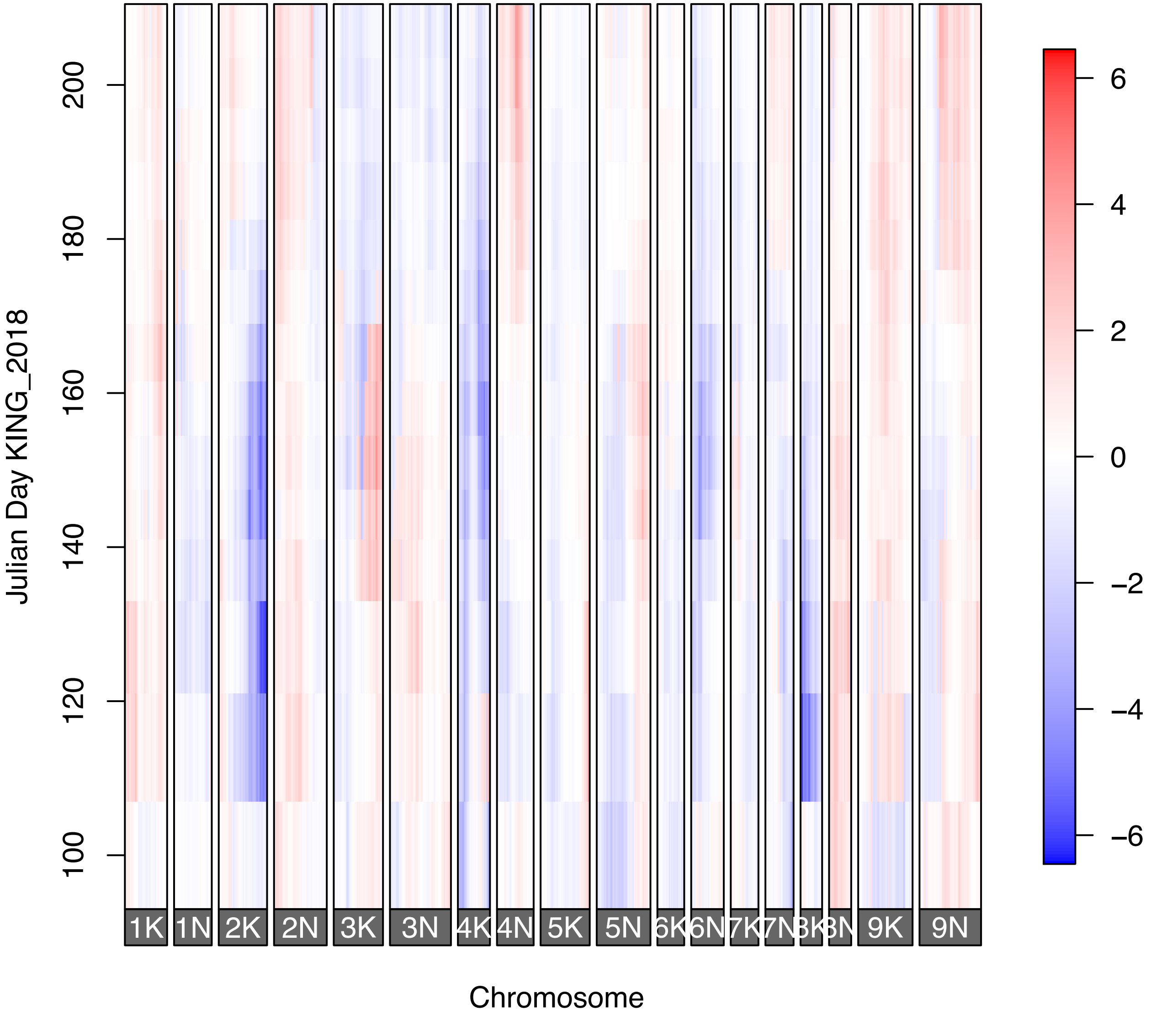
Julian Day KING_2018
Chromosome
1K
1N
2K
2N
3K
3N
4K
4N
5K
5N
6K
6N
7K
7N
8K
8N
9K
9N
100
120
140
160
180
200
6
4
2
0
−2
−4
−6

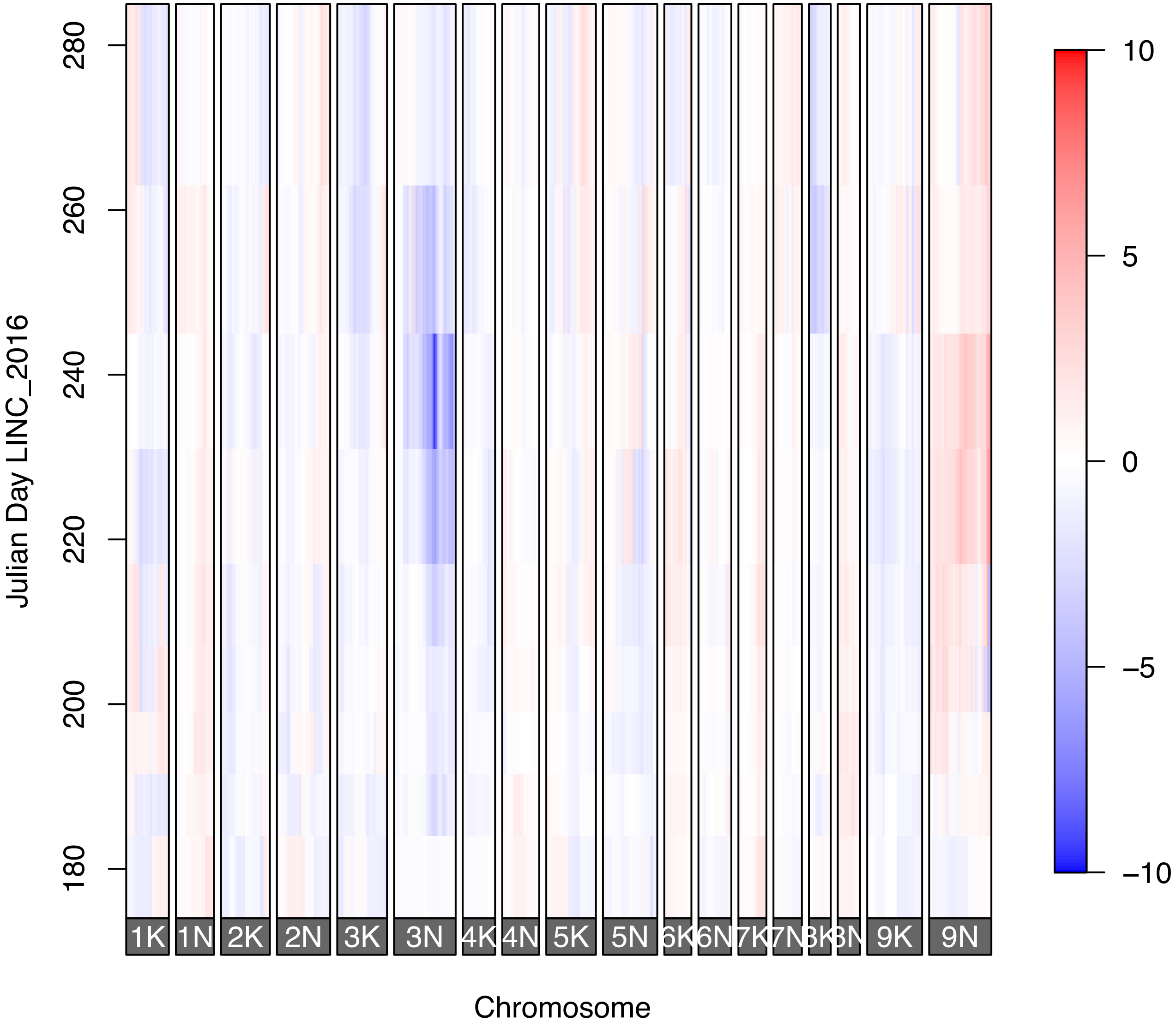
Julian Day LINC_2016
280
260
240
220
200
180
1K
1N
2K
2N
3K
3N
4K
4N
5K
5N
6K
6N
7K
7N
8K
8N
9K
9N
Chromosome
10
5
0
−5
−10

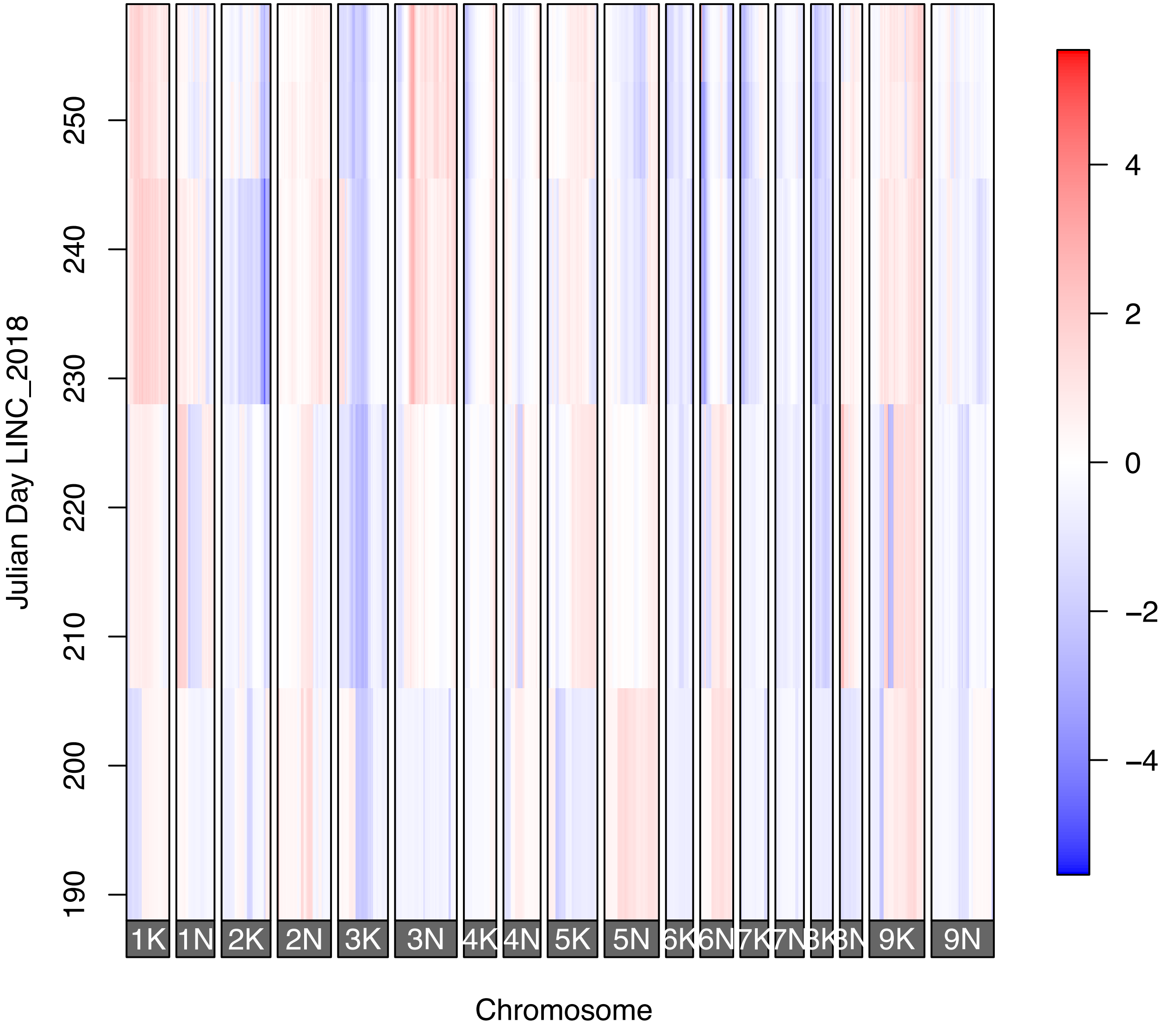

Julian Day LINC_2018
190
200
210
220
230
240
250
1K
1N
2K
2N
3K
3N
4K
4N
5K
5N
6K
6N
7K
7N
8K
8N
9K
9N
Chromosome
4
2
0
−2
−4

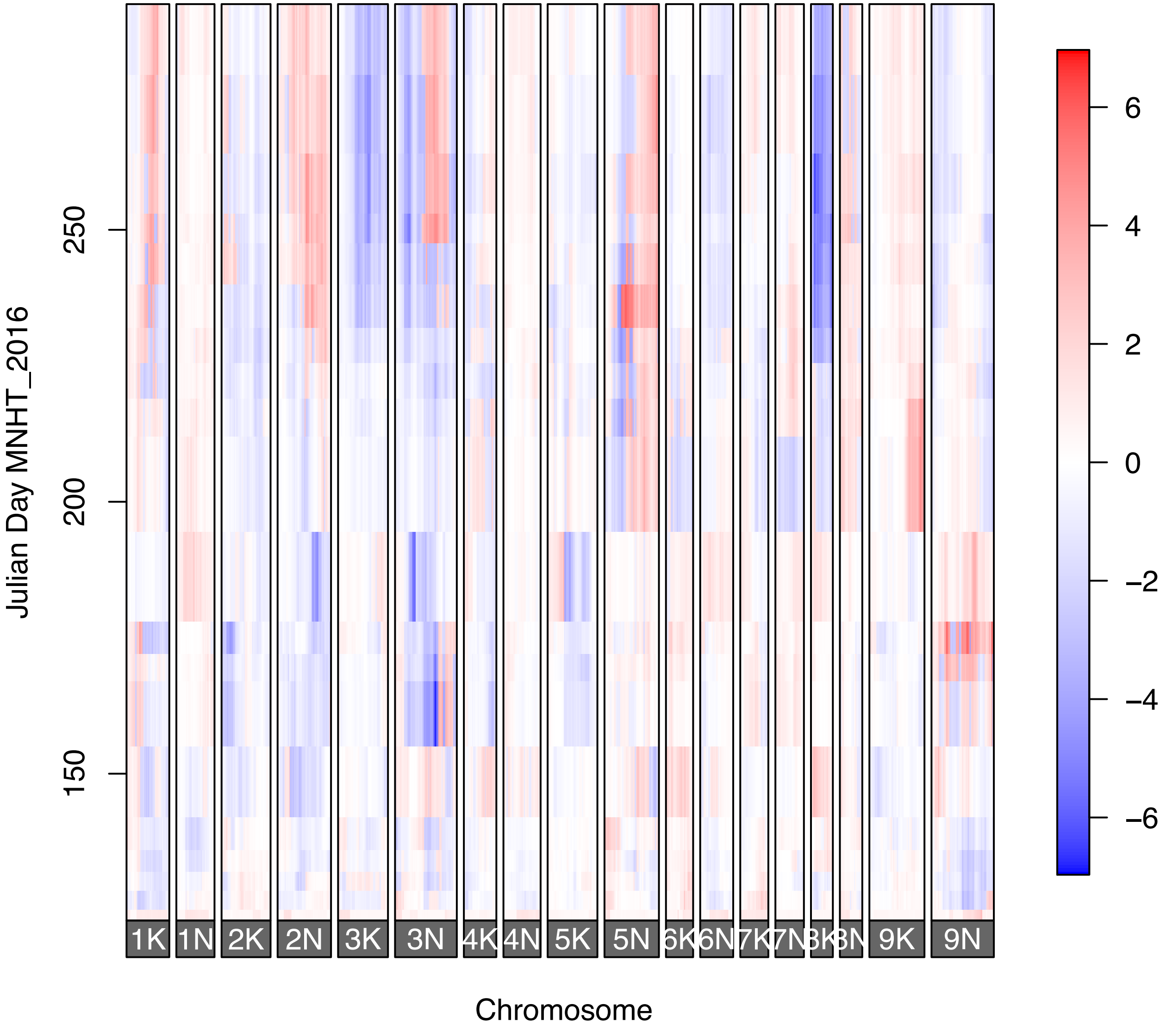
Julian Day MNHT_2016
150
200
250
1K
1N
2K
2N
3K
3N
4K
4N
5K
5N
6K
6N
7K
7N
8K
8N
9K
9N
Chromosome
6
4
2
0
−2
−4
−6

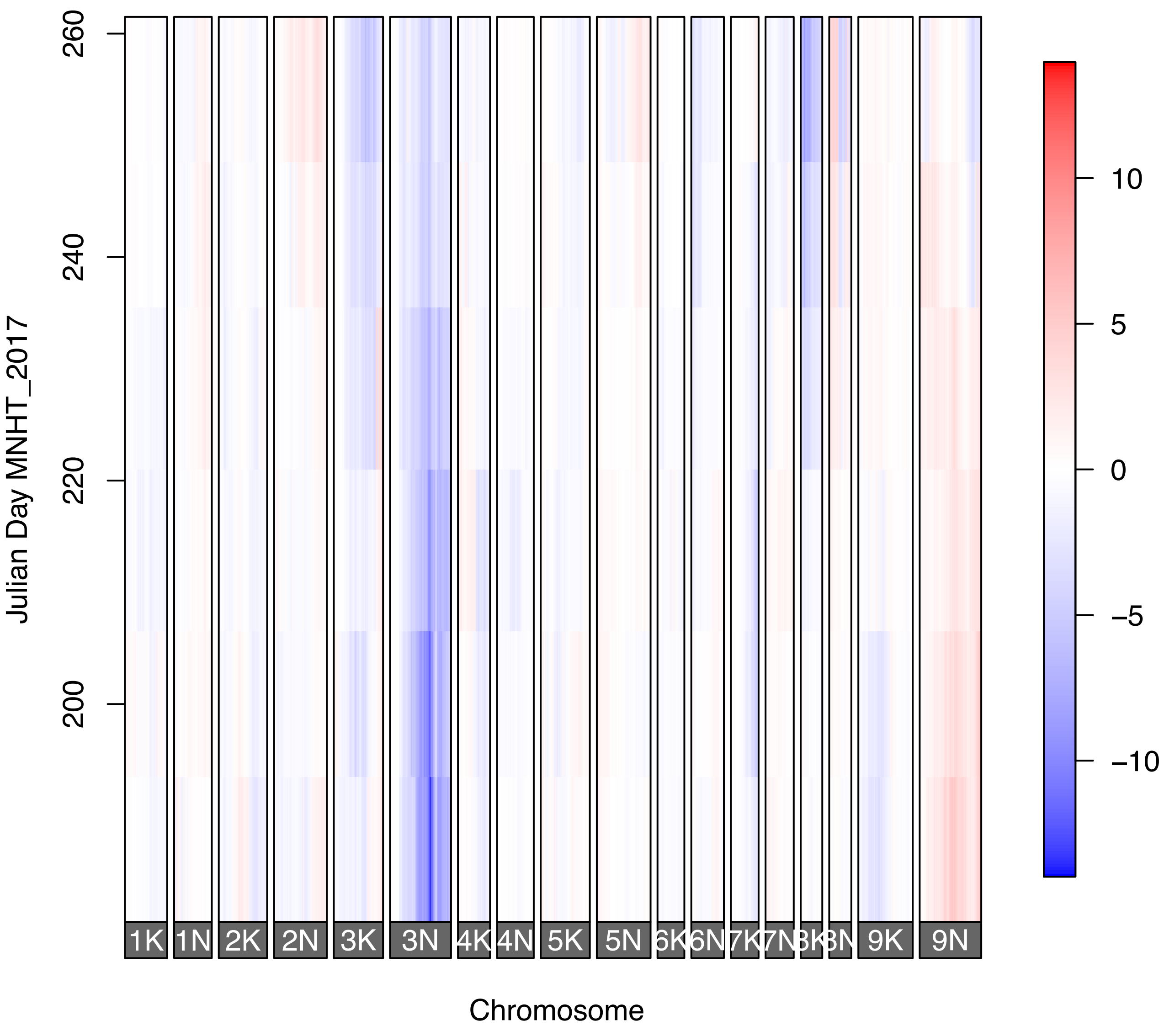
Julian Day MNHT_2017
260
240
220
200
10
5
0
−5
−10
1K
1N
2K
2N
3K
3N
4K
4N
5K
5N
6K
6N
7K
7N
9K
9N
Chromosome

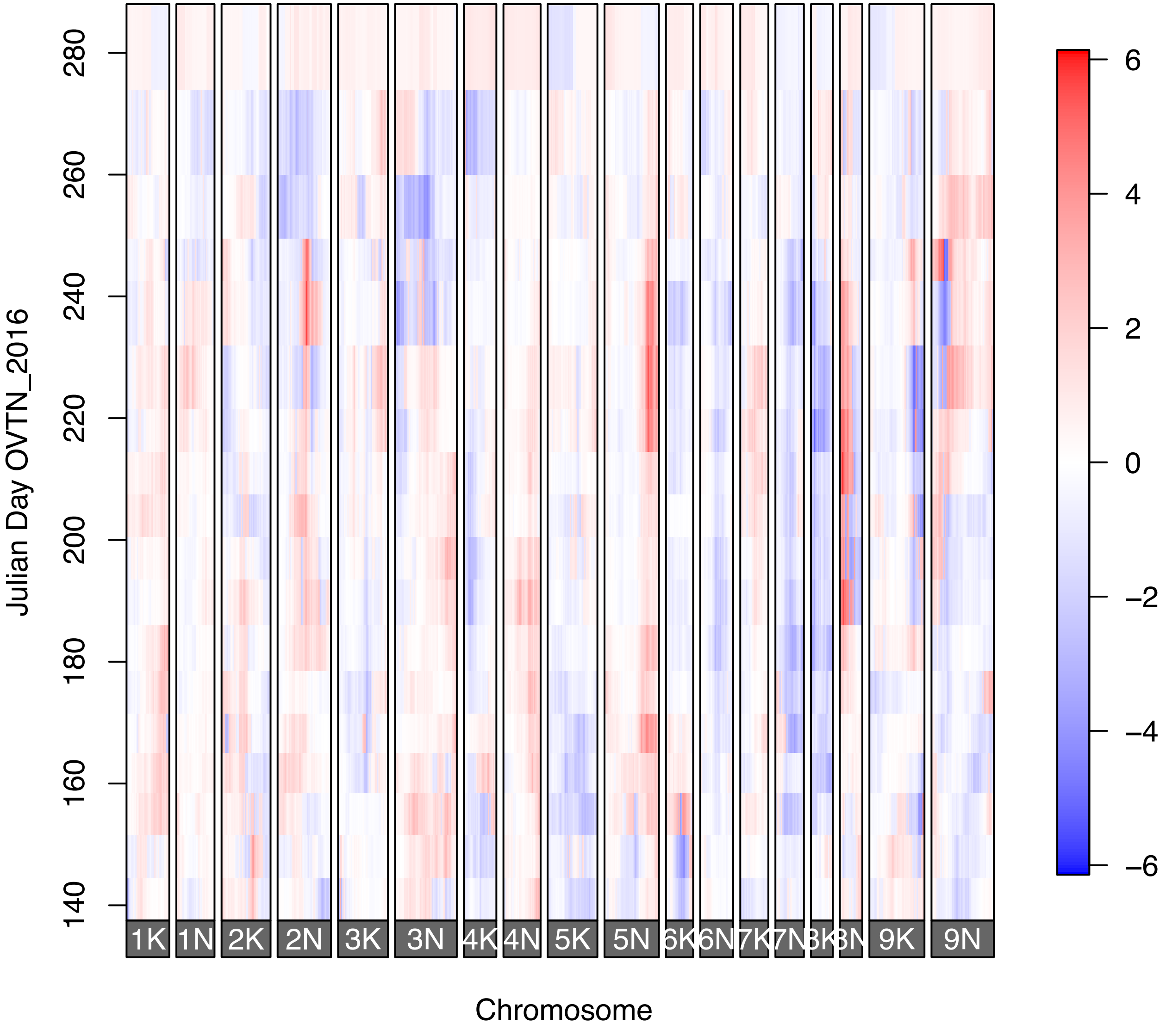
Julian Day OVTN_2016
280
260
240
220
200
180
160
140
1K
1N
2K
2N
3K
3N
4K
4N
5K
5N
6K
6N
7K
7N
9K
9N
Chromosome
6
4
2
0
−2
−4
−6

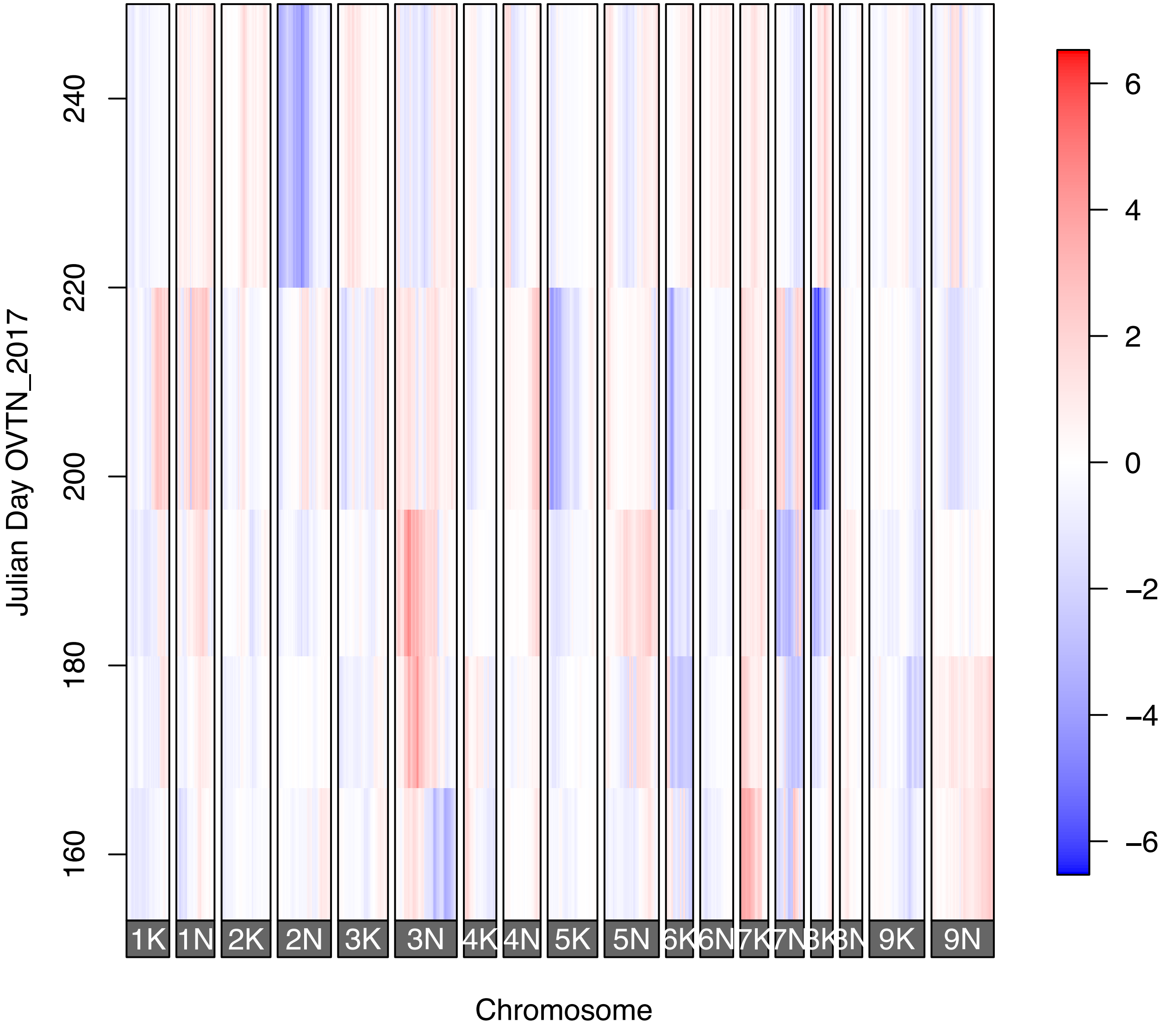

Julian Day OVTN_2017
160
180
200
220
240
Chromosome
1K
1N
2K
2N
3K
3N
4K
4N
5K
5N
6K
6N
7K
7N
8K
8N
9K
9N
6
4
2
0
−2
−4
−6

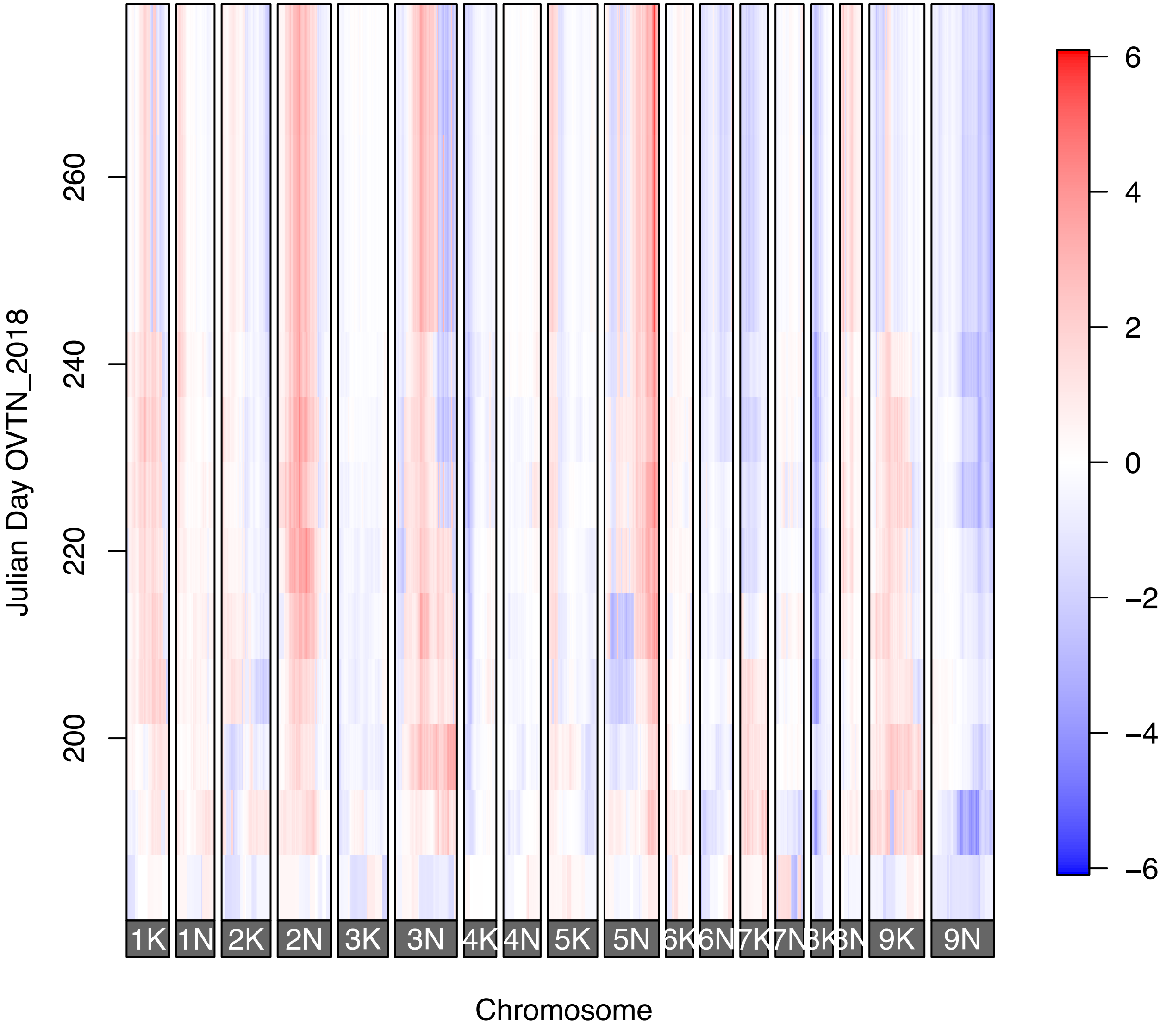

Julian Day OVTN_2018
200
220
240
260
1K
1N
2K
2N
3K
3N
4K
4N
5K
5N
6K
6N
7K
7N
8K
8N
9K
9N
Chromosome
6
4
2
0
−2
−4
−6

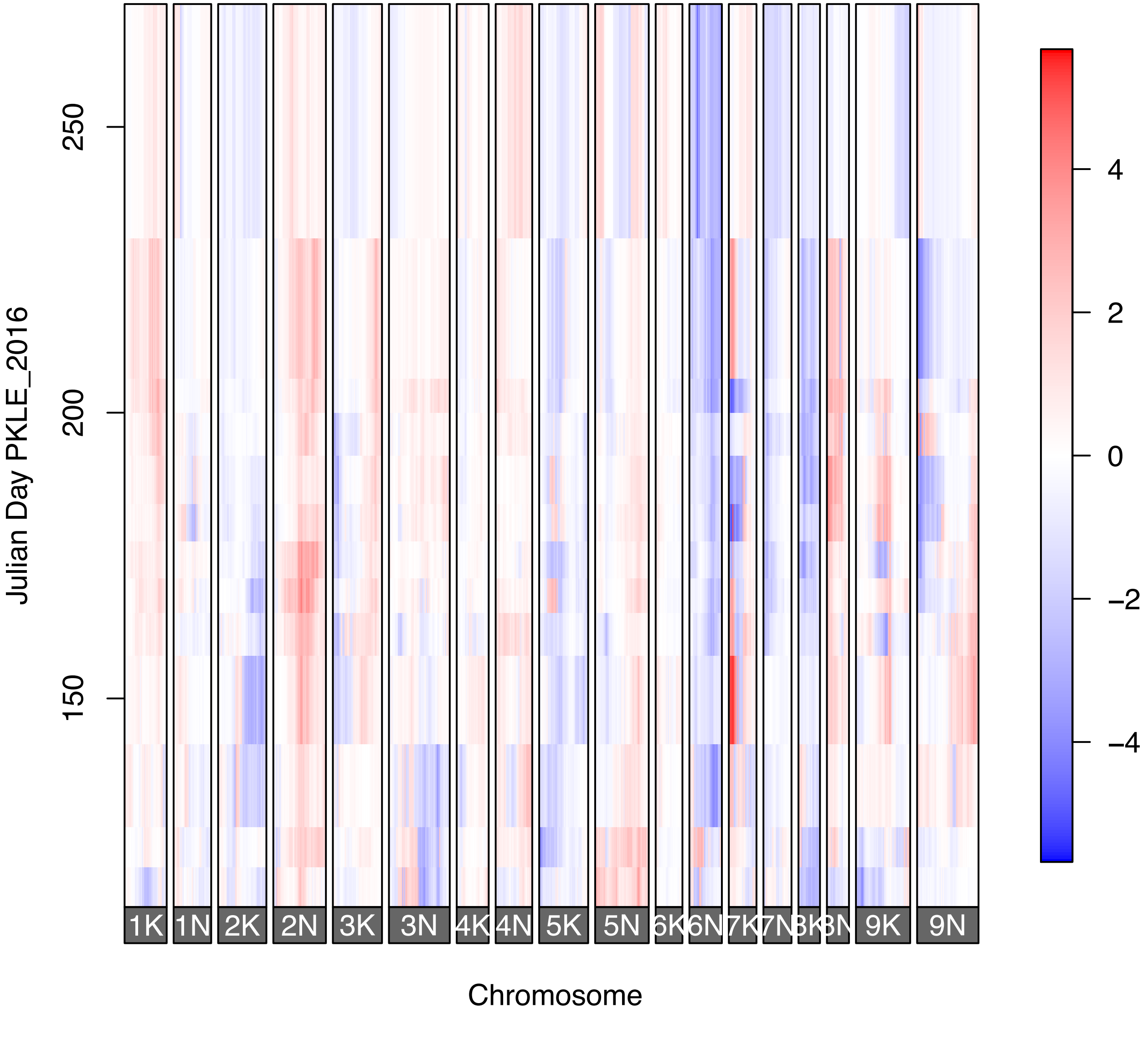

Julian Day PKLE_2016
250
200
150
1K
1N
2K
2N
3K
3N
4K
4N
5K
5N
6K
6N
7K
7N
8K
8N
9K
9N
Chromosome
4
2
0
−2
−4

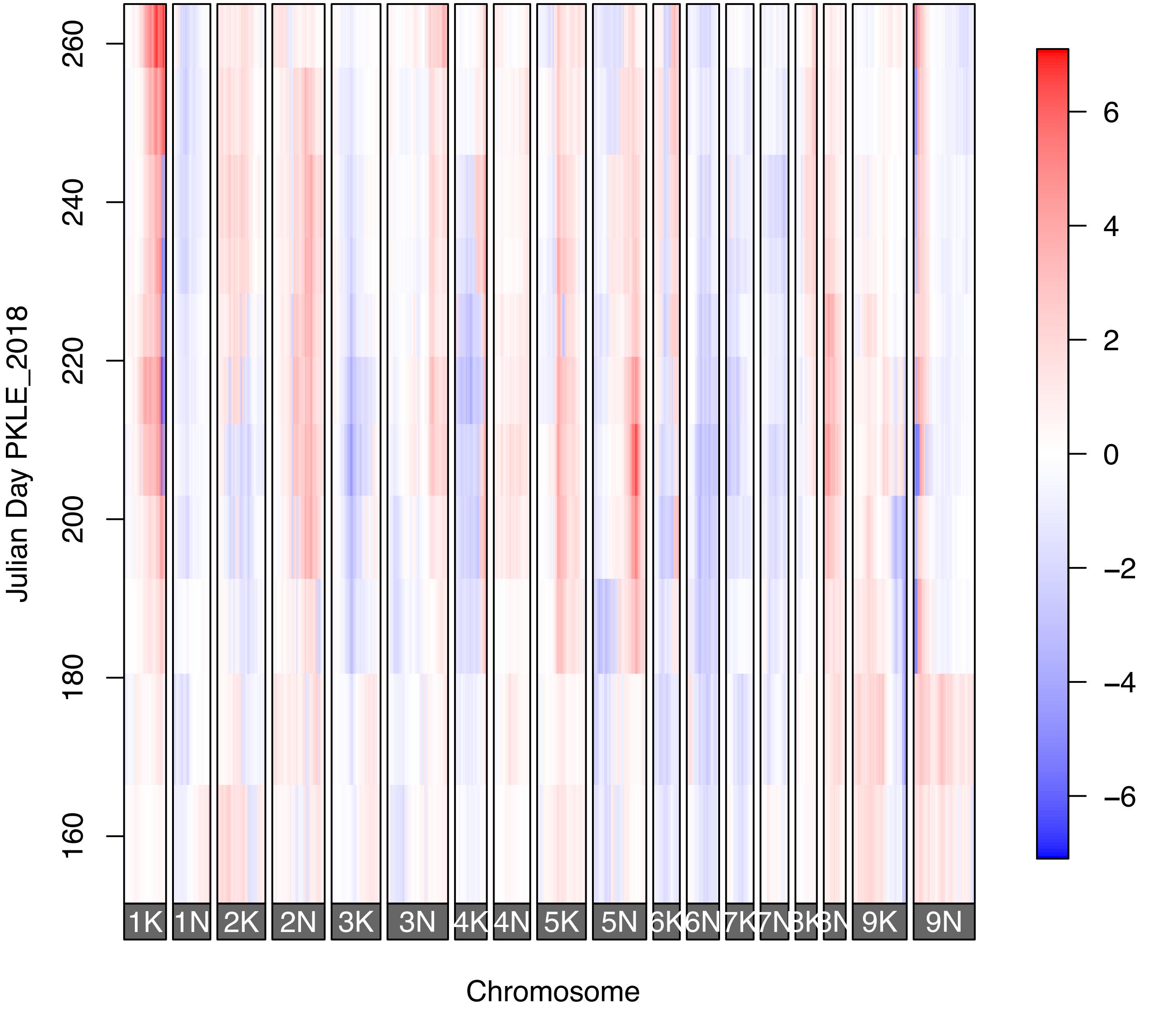
Julian Day PKLE_2018
260
240
220
200
180
160
1K
1N
2K
2N
3K
3N
4K
4N
5K
5N
6K
6N
7K
7N
3K
3N
9K
9N
Chromosome
6
4
2
0
−2
−4
−6

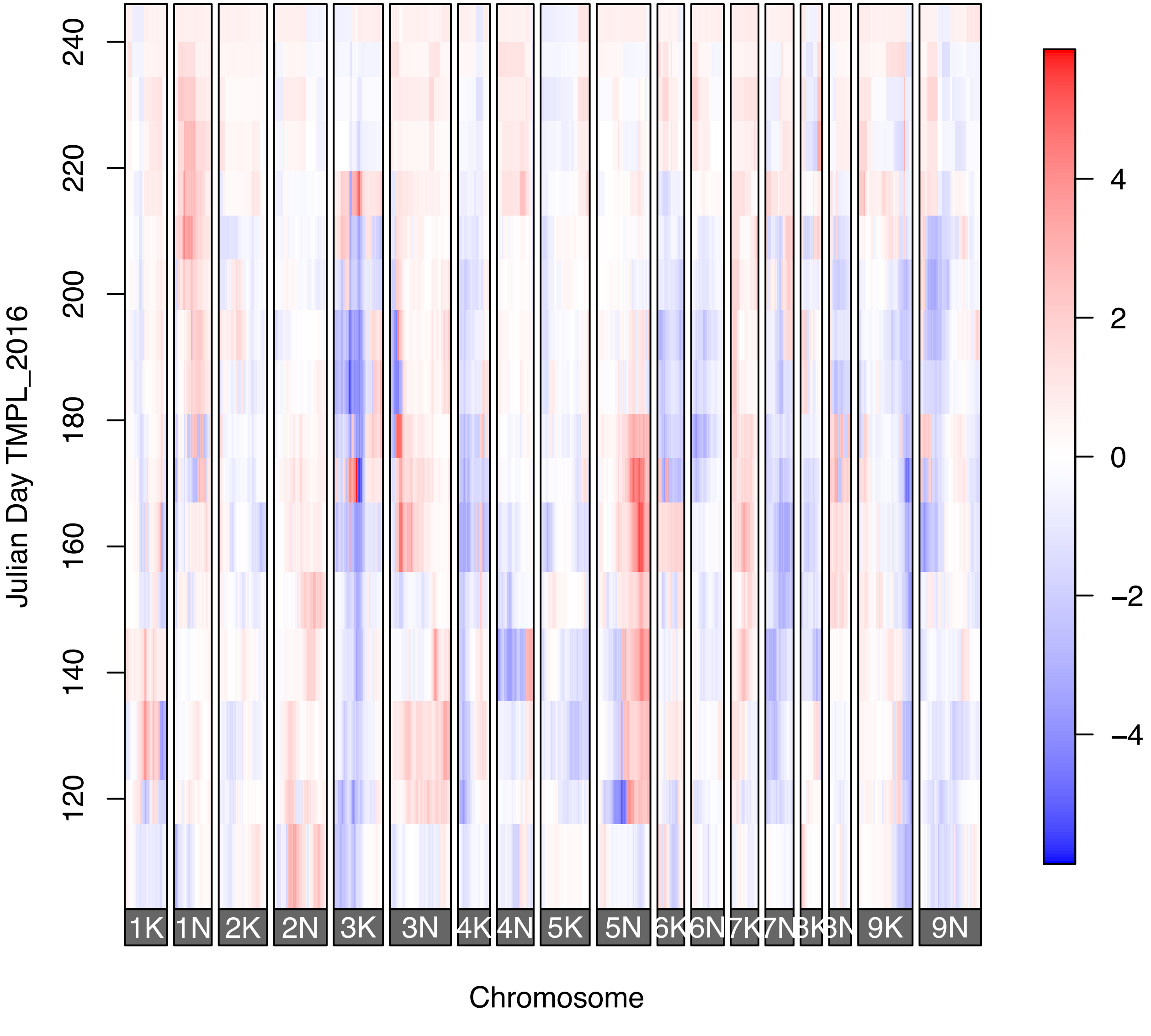

Julian Day TMPL_2016
240
220
200
180
160
140
120
1K
1N
2K
2N
3K
3N
4K
4N
5K
5N
6K
6N
7K
7N
8K
8N
9K
9N
Chromosome
4
2
0
−2
−4

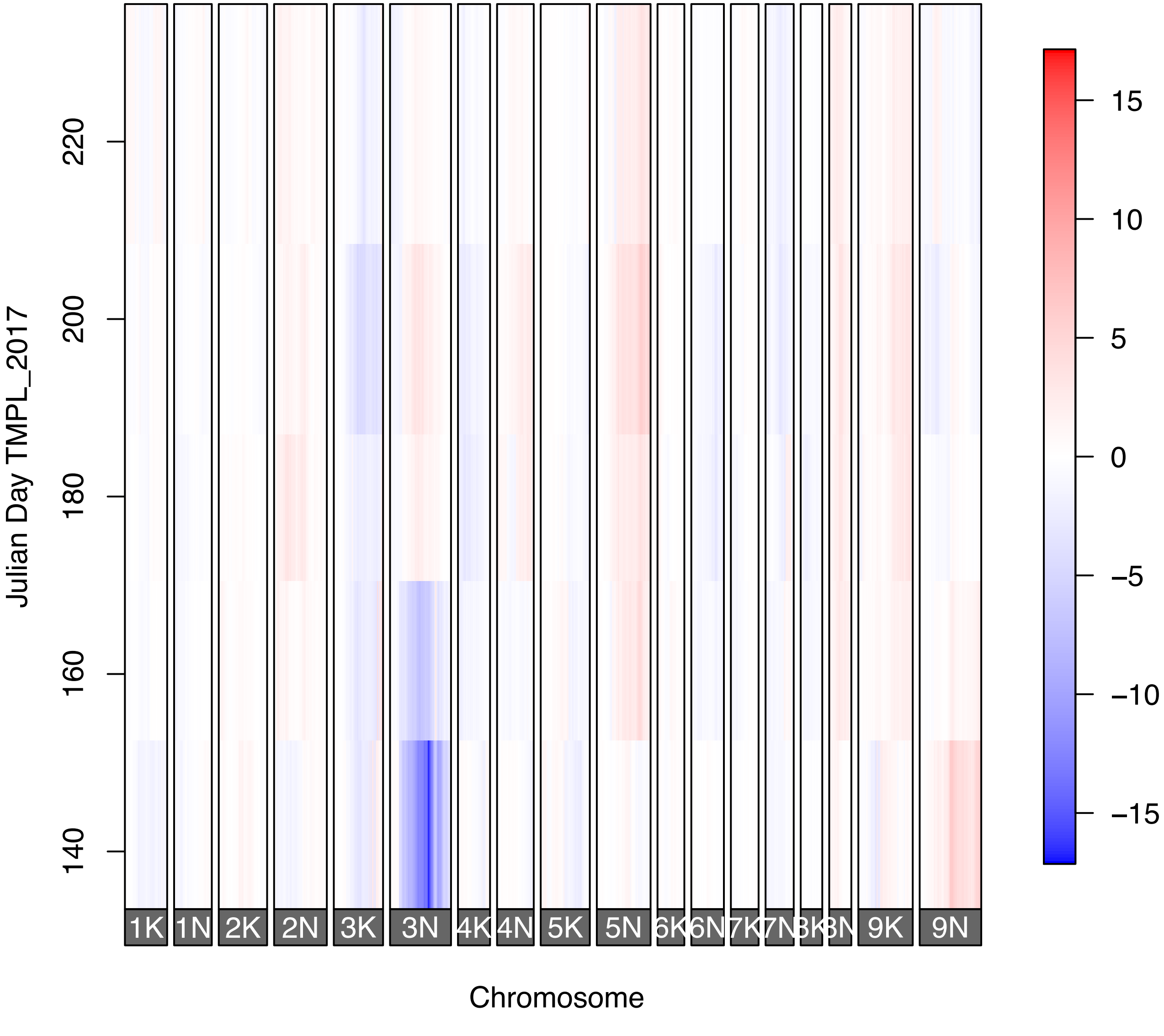

Julian Day TMPL_2017
220
200
180
160
140
1K
1N
2K
2N
3K
3N
4K
4N
5K
5N
6K
6N
7K
7N
8K
8N
9K
9N
Chromosome
15
10
5
0
−5
−10
−15

**Figure S6**: 10% LOD-drop intervals for *Prr1* and *Prr2* (red) overlain on QTLs for morphological and phenological traits. Reproduced with permission from Lowry *et al.*, 2019. Grey boxes highlight *Prr1* and *Prr2*.

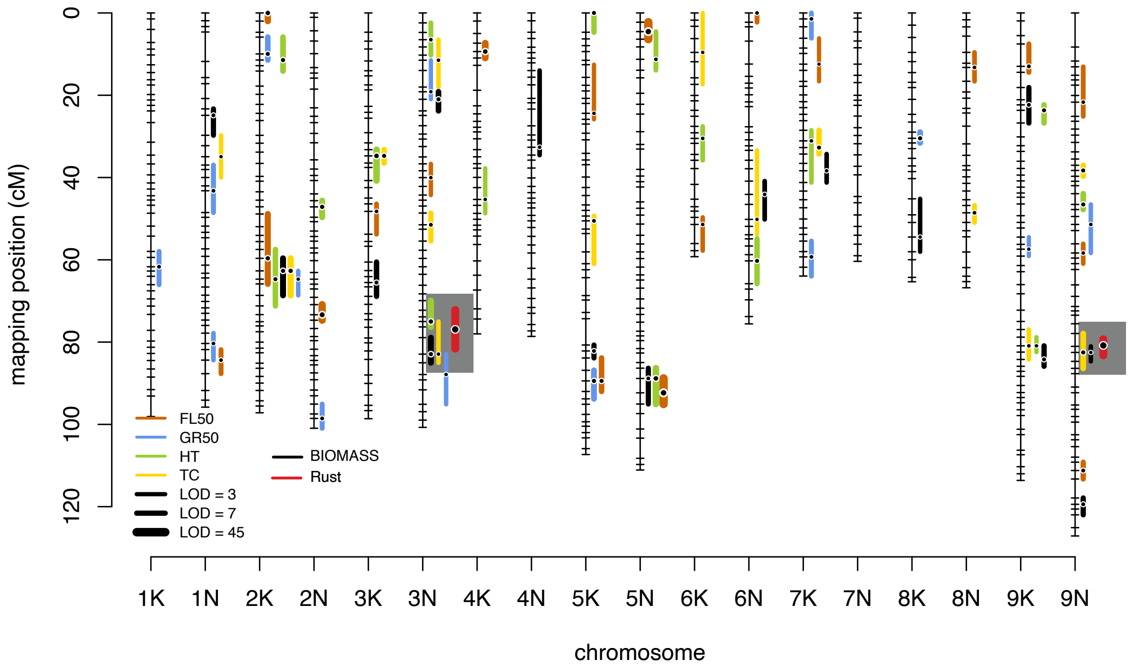

**Figure S6**: 10% LOD-drop intervals for *Prr1* and *Prr2* (red) overlain on QTLs for morphological and phenological traits. Reproduced with permission from Lowry *et al.*, 2019. Grey boxes highlight *Prr1* and *Prr2*.

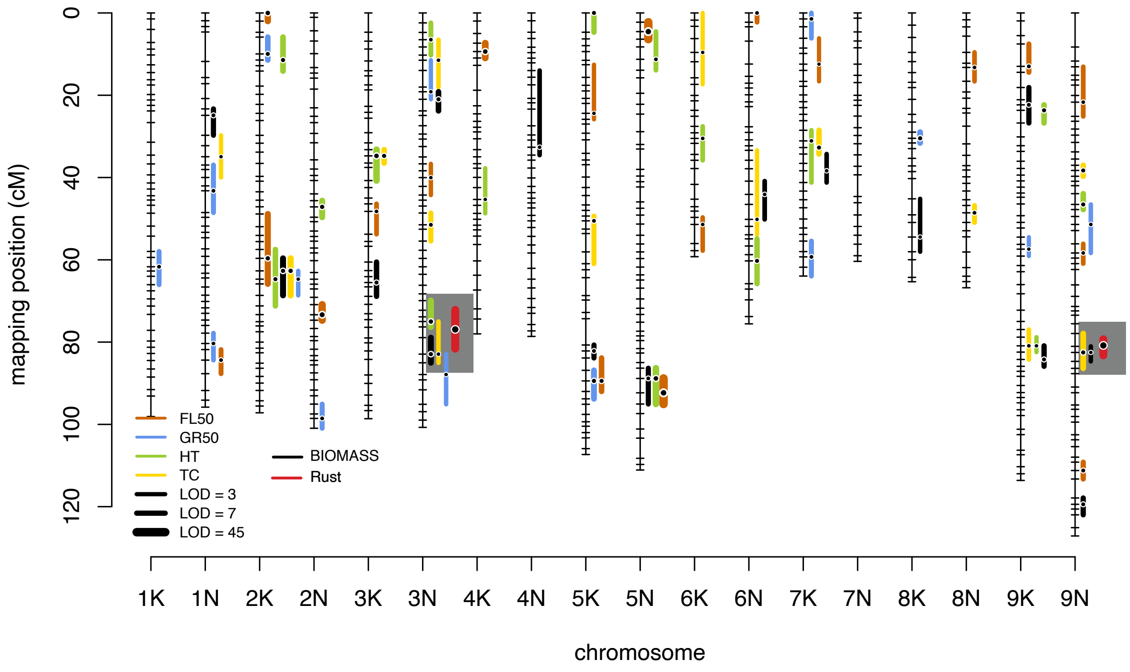

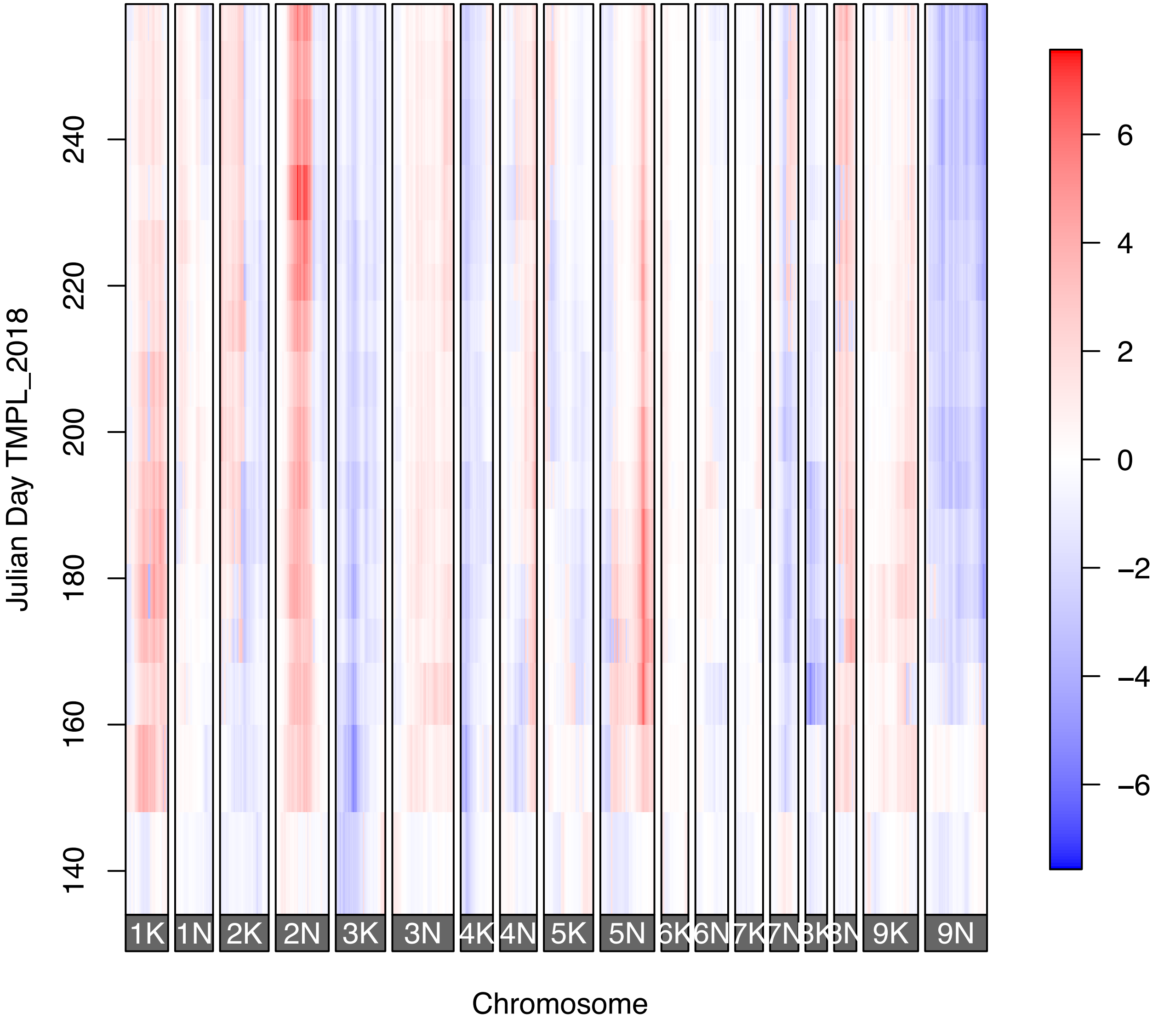

Julian Day TMPL_2018
240
220
200
180
160
140
1K
1N
2K
2N
3K
3N
4K
4N
5K
5N
6K
6N
7K
7N
9K
9N
Chromosome
6
4
2
0
−2
−4
−6
